# Responsiveness to perturbations is a hallmark of transcription factors that maintain cell identity

**DOI:** 10.1101/2020.06.11.147207

**Authors:** Ian A. Mellis, Hailey I. Edelstein, Rachel Truitt, Lauren E. Beck, Orsolya Symmons, Yogesh Goyal, Margaret C. Dunagin, Ricardo A. Linares Saldana, Parisha P. Shah, Wenli Yang, Rajan Jain, Arjun Raj

## Abstract

Our ability to identify the particular transcription factors that maintain cell type is limited. Identification of factors by their cell type-specific expression or their participation in developmental regulation has been only modestly successful. We hypothesized that because cell type is often resilient to perturbations, the transcriptional response to perturbations would identify identity-maintaining factors. We developed Perturbation Panel Profiling (P^3^) as a framework for perturbing cells in dozens of conditions and measuring gene expression responsiveness transcriptome-wide. Applying P^3^ to human iPSC-derived cardiac myocytes showed that transcription factors known to function in cardiac differentiation and maintenance were among the most frequently up-regulated (most responsive). We reasoned that one potential function of responsive genes may be to maintain cellular identity. We identified responsive transcription factors in fibroblasts using P^3^ and found that suppressing their expression led to enhanced reprogramming efficiency. We propose that responsiveness to perturbations is a property of factors that help maintain cellular identity.

## Introduction

Cells of a particular cell type likely maintain many of their specific behavioral properties across a range of conditions. The maintenance of these properties can be the result of the activity of certain transcription factors, and the identification of these factors may be useful for manipulating cellular identity (*1*). However, it has proven difficult to predict which transcription factors, of the many transcription factors expressed in a given cell type, are responsible for the regulation of that cellular identity.

Transcription factors can be involved in cellular identity in a number of ways. Two key functional properties of transcription factors are their ability to be *identity-establishing* or *identity-maintaining*, meaning that they are able to control the expression of genes that will establish or maintain the characteristics of a particular cell type, respectively. A correlative property of a transcription factor is its cell type-specificity, meaning the degree to which its expression is almost entirely restricted to that one cell type and not to others. Cell type-specificity is a relatively easy property to measure, given the ubiquity of large gene expression datasets. Several recent approaches have mined these large gene expression datasets to try and identify functionally important transcription factors based on their cell type-specificity or inferred activity in particular tissues (*2*–*6*). The cell type-specificity of a transcription factor, however, is fundamentally correlative and does not *per se* yield any information about the functional properties of identity establishment or maintenance, and so the pool of identified transcription factors may contain several non-functional candidates. Using cell type-specificity to identify causative factors can also suffer from the converse problem: it could be that there are factors that can establish or maintain cell type but are not cell-type specific in their expression or activity. Such factors would also not be easily identified by methods relying on cell type-specificity.

A related approach to identifying cellular identity-establishing or -maintaining factors is to focus on factors known to be involved in the developmental process responsible for differentiating cells into a particular type (*7*–*14*). However, in other contexts, such as transdifferentiation of cells grown in culture, there may be factors that can establish cell type via trajectories that do not mirror normal development, which could be missed by those approaches (*15*, *16*).

We hypothesized that the transcriptional response to perturbations would help identify genes whose expression is important for maintaining cell type. We envisioned two scenarios with distinct transcriptional response profiles to help maintain cellular identity. In the first scenario, critical transcription factors maintain stable expression in the face of perturbations, that is, they are the steady, non-responsive “pillars” of expression. In the second scenario, critical transcription factors are highly reactive and frequently respond to perturbations to ensure that cells maintain their type; that is, they are hyper-responsive “springs” of expression; **Fig. 1A**. Transcriptionally profiling populations of cells subjected to a large panel of perturbations would allow us to discern whether one scenario was favored or if there was equipoise between the scenarios. Other perturbation profiling datasets have been generated, but their experimental designs have been geared towards other goals like drug target identification (*17*–*19*).

**Figure 1:**
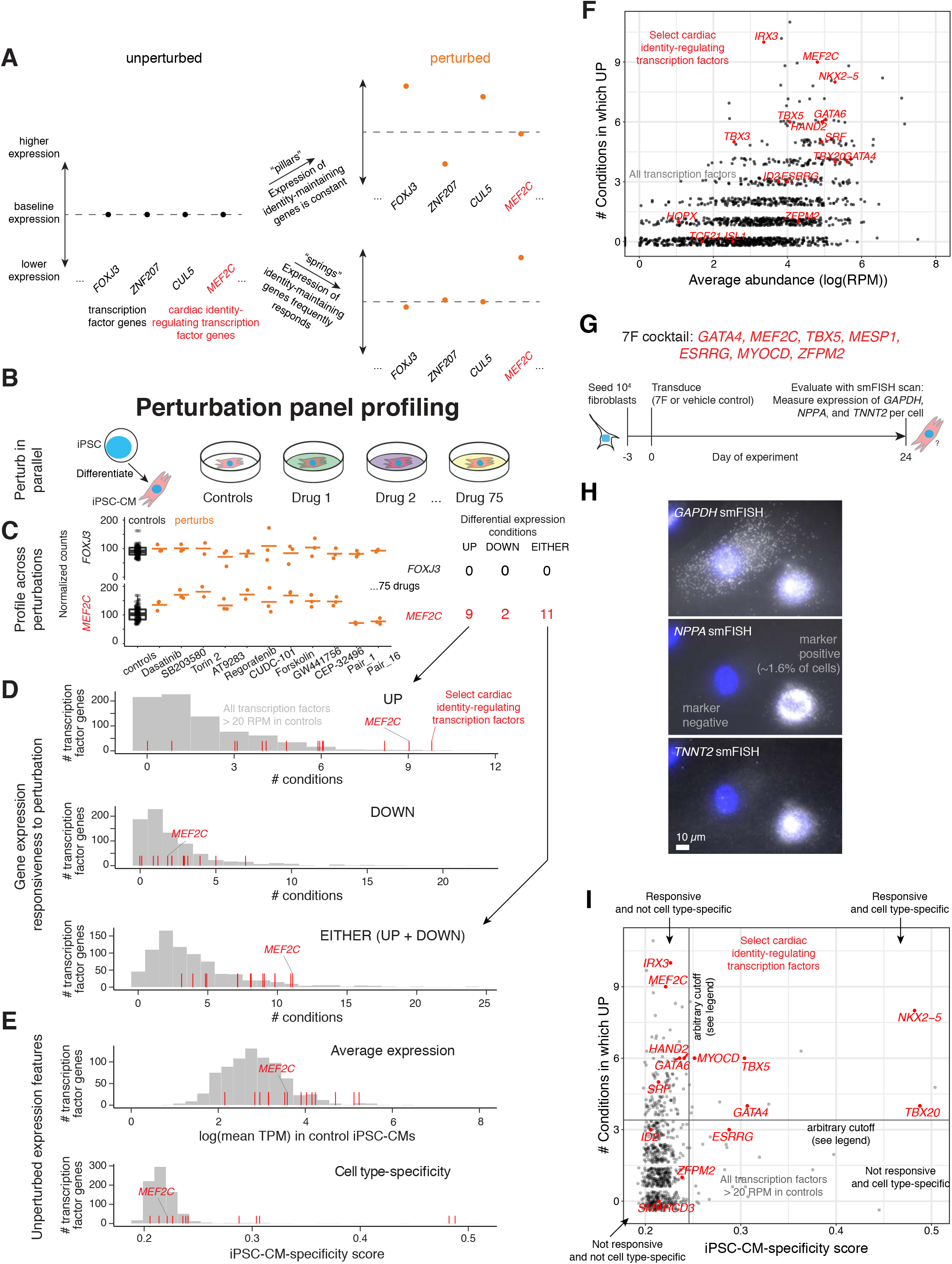
Perturbation panel profiling of iPSC-derived cardiac myocytes. A) Conceptual guide for different perturbation-response models of identity-maintaining genes. In the “pillars” model, identity-maintaining genes are relatively unresponsive to perturbation, whereas non-identity-related genes do respond. In the “springs” model, identity-maintaining genes are more often responsive to perturbation than other genes. B) Perturbation panel profiling experiments and analysis: parallel cultures of human iPSC-derived cardiac myocytes (iPSC-CM), exposed to 75 different perturbation conditions in triplicate (see Table S1) or vehicle control. C) Parallel RNA extraction and library preparation for RNAtag-seq of perturbed and control samples followed by differential expression analysis. Per gene, differential expression in each condition relative to vehicle control samples followed by quantification of the number of conditions in which differential expression was observed. Representative results for two genes, *FOXJ3* and *MEF2C*, are shown. D) Responsiveness to perturbation analysis of all expressed transcription factor genes. All transcription factor genes expressed at 20 RPM or greater in controls were considered (grey, histogram). Pre-registered set of cardiac myocyte identity-establishing and -maintaining genes shown in red. For each gene, the number of conditions in which any differential expression (top) differential expression down (middle) or up (bottom). E) Static expression feature analysis of all expressed transcription factor genes. All transcription factor genes expressed at 20 RPM or greater in controls were considered (grey, histogram). iPSC-derived cardiac myocyte-specificity score is Jensen-Shannon specificity to iPSC-derived cardiac myocytes without GTEx Skin and Heart (See Methods). Pre-registered set of cardiac myocyte identity-regulating genes shown in red. For each gene, the average expression level in vehicle control samples (top) or iPSC-derived cardiac myocyte-specificity score (bottom, see Methods). F) Comparison of responsiveness (number of iPSC-CM conditions in which up-regulated) and mean expression (mean RPM in control iPSC-CMs) for all expressed transcription factor genes. Vertical-only jitter added for visualization of all points with the same responsiveness value. G) Comparison of responsiveness and iPSC-derived cardiac myocyte-specificity scores for all expressed transcription factor genes. Vertical-only jitter added for visualization of all points with the same responsiveness value. Quadrant-separating lines (“arbitrary cutoffs”) chosen as elbows of the cumulative distribution function of each feature over all transcription factor genes expressed at 20 RPM or greater in controls. H) Schematic of cardiac transdifferentiation experimental workflow. See Methods for full details. I) Representative images of *GAPDH*, *NPPA*, and *TNNT2* smFISH of 7F-transduced fibroblasts after 24 days, demonstrating a cell that expresses marker genes *NPPA* and *TNNT2* at high levels and one cell that does not. Cells called “marker-positive” if *TNNT2* spots > 19 or *NPPA* spots > 24, thresholds chosen arbitrarily. 0 cells in negative control MXs-empty condition are “marker-positive”. Scale bars are 10 μm.

Here, we developed an experimental and analytical pipeline, perturbation panel profiling (P^3^), to measure the genome-wide transcriptional responses to a large set of perturbations in human iPSC-derived cardiac myocytes and fibroblasts. We demonstrated that well-known regulators of cardiogenesis are amongst the most responsive in their expression to a wide variety of perturbations in cardiac myocytes; i.e., they act as springs rather than pillars. We then applied this same protocol to fibroblasts to identify highly responsive genes in that cell type. We showed that lowering the expression of fibroblast-responsive factors identified by P^3^ led to an enhancement in the cells’ ability to reprogram into induced pluripotent stem cells. This enhancement suggested an increased pliability of cell type; i.e., suppression of responsive transcription factors reduced fibroblast identity maintenance. In sum, our results demonstrate that responsiveness to perturbation may represent a property of transcription factor genes that are important for cell type maintenance.

## Results

### Transcription factor genes that regulate cellular identity are upregulated in cardiac myocytes after many different perturbations

To determine whether genes that are critical for cardiogenesis are more or less often responsive to perturbation than other similarly highly expressed genes, we designed an experimental and analytical workflow, P^3^. P^3^ consists of growing highly pure cell type populations, exposing them to a panel of dozens of different drugs in parallel, and performing RNAtag-seq for transcriptome-wide gene expression profiling in each condition. We then measure changes in gene expression across all conditions. We chose to initially apply P^3^ to human iPSC-derived cardiac myocytes (iPSC-CMs) because of the ease of culture, applicability to emerging therapies, and well-established developmental trajectories. To identify potential factors in an unbiased way, we sought to perturb cells such that, in aggregate, we affected as much of the transcriptome as possible. Therefore, we sought to maximize the number of signaling cascades we perturbed, identifying 100 small molecules from the SelleckChem Bioactive library targeting any kinases or G protein-coupled receptors (GPCR) and minimizing overlap in their annotated targets (**Table S1**). We administered these drugs individually or in pairs, for a total of 75 different perturbation conditions (**Fig. S1A**).

We cultured hundreds of parallel bulk samples of iPSC-derived cardiac myocytes, cultured under standard (i.e., control, n = 63 replicate samples, spread evenly across all plates) and different (N=75) perturbed conditions (**Fig. 1B, Fig. S1A,** n= 3 replicate samples per condition). The dose for each drug was chosen to be 100 times the median IC50 values (or 10 times for each drug in a pair) culled from the literature and product materials (we determined that 10x - 100x were the appropriate dose factor empirically; data not shown; see Methods), and ultimately applied each drug at one drug-specific dose in triplicate to samples that we further analyzed with RNAtag-seq (*20*). After treating iPSC-derived cardiac myocytes with each compound or DMSO-only exposure for 4 days, we found that 63 out of 75 conditions had wells with beating cells similar in morphology to vehicle-treated cells and without evident cell death (**Fig. S1B,C**).

We next conducted transcriptome-wide gene expression profiling of the hundreds of perturbed (and control) samples of cardiac myocytes **(Fig. 1C)**. We performed RNAtag-seq in batches of 96 samples, distributing DMSO controls evenly across each batch. Gene expression profiles across the 458 quality-controlled samples (454 excluding 4 HeLa samples used as an outgroup) clustered first by cell type, and then often by perturbation condition (**Fig. S2**). We identified hundreds to thousands (0 - 6862 genes per condition) of differentially expressed genes in most perturbed conditions relative to corresponding cell type controls (**Fig. S3A**, Methods). We sequenced our libraries to a depth at which we estimated that fully doubling the number of reads for all 458 samples would have increased the number of detected differentially expressed genes by at most only 48% for genes expressed at an average of 20 RPM or greater in controls (note that sequencing to half the depth would have reduced the number of detected differentially expressed genes by 29%; **Fig. S4**).

In order to ensure our analysis was sufficiently unbiased, we verified that the majority of the coding genome was affected by a subset of our 60+ perturbations. We quantified the number of perturbations which resulted in differential expression in iPSC-derived cardiac myocytes for each gene **(Fig. 1C)**, limiting our analyses to genes which were expressed at 20 RPM or higher in control iPSC-derived cardiac myocytes. We found that 94.5% of genes expressed at 20 RPM or greater in iPSC-derived cardiac myocytes were differentially expressed in at least 1 perturbation condition relative to the cell type controls (**Fig. S3**), demonstrating that the compounds in our library were able to perturb the vast majority of the transcriptome. We focused our subsequent analyses on transcription factors, as they control the expression of genes relevant to cellular identity and homeostasis.

Next, we determined the degree to which individual transcription factors were responsive in the iPSC-derived cardiac myocyte dataset. We quantified the number of perturbations of iPSC-derived cardiac myocytes that resulted in differential expression (adj. p-value < 0.1) of a transcription factor versus controls. In addition, we wanted to ensure that responsiveness did not merely identify genes which were highly expressed or that only highly expressed genes were detectably responsive. We found that expression level did not entirely correlate with detected responsiveness: many lowly expressed transcription factors were responsive to perturbations and many highly expressed transcription factors were not responsive to perturbation. However, we observed that multiple critical cardiac transcription factors frequently responded to perturbations, including *MEF2C, NKX2-5, GATA4, HAND2, TBX5*. **(Fig. 1D-F)**. Furthermore, gene ontology analysis of all responsive genes (not just transcription factors) identified a clear over-enrichment for genes related to cardiac function (e.g., structural components such as sarcomeres, myofibrils) relative to all highly expressed genes. By contrast, non-responsive genes did not demonstrate this over-enrichment (**Fig. S6**). Prominently among the transcription factors were 6 of the 7 members of the “7F” cocktail, whose combined expression has previously been reported to induce transdifferentiation of fibroblasts into cardiac myocytes (*11*).

We wanted to confirm that these factors could enable such transdifferentiation in the specific fibroblast line we originally used to generate the iPSC-derived cardiac myocytes, so we overexpressed them in isogenic GM00942 fibroblasts by viral transduction. Using single-molecule RNA FISH (*21*), we found that around 1% of the cells expressed the cardiac myocyte-specific marker genes *TNNT2* and *NPPA*. This efficiency is comparable to previously reported results (*11*, *22*). The appearance of these markers suggested that these cells were at least partially reprogrammed to a cardiac myocyte-like state (**Fig. 1H,I, S5A-E**), a finding we confirmed in two other fibroblast lines (**Fig. S5F**). These transdifferentiation results confirm the importance of the 7F factors in regulating cardiac myocyte identity. Notably, 6 of these 7 factors were among the top hits in our P^3^ experiments. Thus, transcription factors critical for regulation of cardiac myocyte identity and cardiac function appear to be “springs”, i.e., frequently responsive to perturbation, rather than “pillars” that hold steady in the face of perturbation. There are many reasons why these transcription factors might be more responsive to perturbations, including that they are part of a homeostatic response to perturbation or that they are simply downstream targets of many signaling pathways in the cells whose identity they regulate (see Discussion). One possible model for responsiveness is that factors important for identity maintenance would frequently upregulate to help preserve identity in the face of a perturbation. A prediction of this model is that high responsiveness could signify that a transcription factor is important for maintenance of cellular identity. In support of this model, several of these factors have been demonstrated to be identity-maintaining transcription factors, particularly NKX2-5, GATA4, and TBX5 (*23*–*25*). It is of course important to note that responsiveness may also correlate with a number of other properties that a factor may have (*26*).

Responsiveness is a property of a gene’s transcriptional regulation that appeared to be associated with identity-regulating factors. Most approaches to date have, however, focused on cell type-specificity, meaning whether expression of a gene is specific to a given cell type or tissue type. To compare responsiveness to cell type-specificity, we used a cell type-specificity score, which is a metric that combines expression levels and cell type-specificity as defined by various publicly available datasets. We calculated cell type (or tissue)-specificity scores for all genes in each tissue included in the GTEx dataset based on average expression levels (See Methods, (*27*, *28*)). We replaced the heart and skin GTEx samples with expression data generated from our control iPSC-derived cardiac myocytes and GM00942 fibroblasts, respectively, in order to most directly compare expression features from P^3^ experiments with cell type-specificity scores. We compared the responsiveness of factors identified in our iPSC-derived cardiac myocyte and GM00942 fibroblast P^3^ experiments to the iPSC-derived cardiac myocyte- or GM00942 fibroblast-specificity score, respectively **(Figures 1E,G).** We found that responsiveness and cell type-specificity score did not equate (similar results held when using all GTEx samples, **Fig. S7**). Of 14 select factors known to be relevant to cardiac myocyte biology, we found that cell type-specificity score only identified 5 of them, whereas responsiveness identified 10. However, cell type-specificity was far more selective, in that of the top 24 cell type-specific factors, 5 belonged to the 14 known factors, whereas responsiveness identified 123 factors in addition to the 10 of the 14 known ones. It is possible that these additional factors represent further identity-maintaining factors, or that they are merely other responsive factors with no functional role in cellular identity regulation. Further studies will be required to make such a determination (see Discussion).

### Knockdown of putative identity-maintaining factors identified by P^3^ enhances reprogramming

The data generated from applying P^3^ to iPSC-CMs suggested that genes which maintain identity are springs (frequently responsive to perturbations), as opposed to pillars. Specifically, the experiments demonstrated that responsiveness identified a cadre of genes known to, in part, regulate maintenance. Hence we reasoned that responsiveness might also identify previously undescribed factors that maintain cellular identity. One way to test this hypothesis is in reprogramming from a source to target cell type: we reasoned that the downregulation of factors that are responsive in the source cell type may interfere with source cell type maintenance, manifesting as an enhancement of reprogramming to the target cell type. We tested this possibility by focusing on the induction of induced pluripotent stem cells from fibroblasts, i.e., fibroblast reprogramming.

To test the hypothesis that P^3^ can reveal genes that, if removed, would inhibit identity-maintenance, we first performed P^3^ on isogenic dermal fibroblasts (GM00942, Coriell, **Fig. 2A**) to find responsive candidate transcription factor genes which we could then evaluate for identity-maintaining behavior. Similar to our iPSC-derived cardiac myocyte studies, we found that 62 out of 75 fibroblast culture conditions had wells with cells with fibroblast morphology and without evident cell death. We also observed similar numbers of detectably differentially expressed genes in fibroblasts as in iPSC-derived cardiac myocytes across the perturbation panel (**Fig. S3**). Responsive genes identified in our P^3^ analysis again did not equate to highly expressed or cell type-specific factors in a parallel analysis to that described above for iPSC-CMs (**Fig. S8A,B**, **S9**). Further, Gene Ontology analysis on all responsive genes (not just transcription factors) revealed an over-enrichment for adhesion-related cellular components (e.g., cell-substrate junction) that are likely related to fibroblast function (**Fig. S6**). This enrichment mirrors the enrichment of cardiac myocyte components amongst the set of responsive genes in iPSC-derived cardiac myocytes.

**Figure 2:**
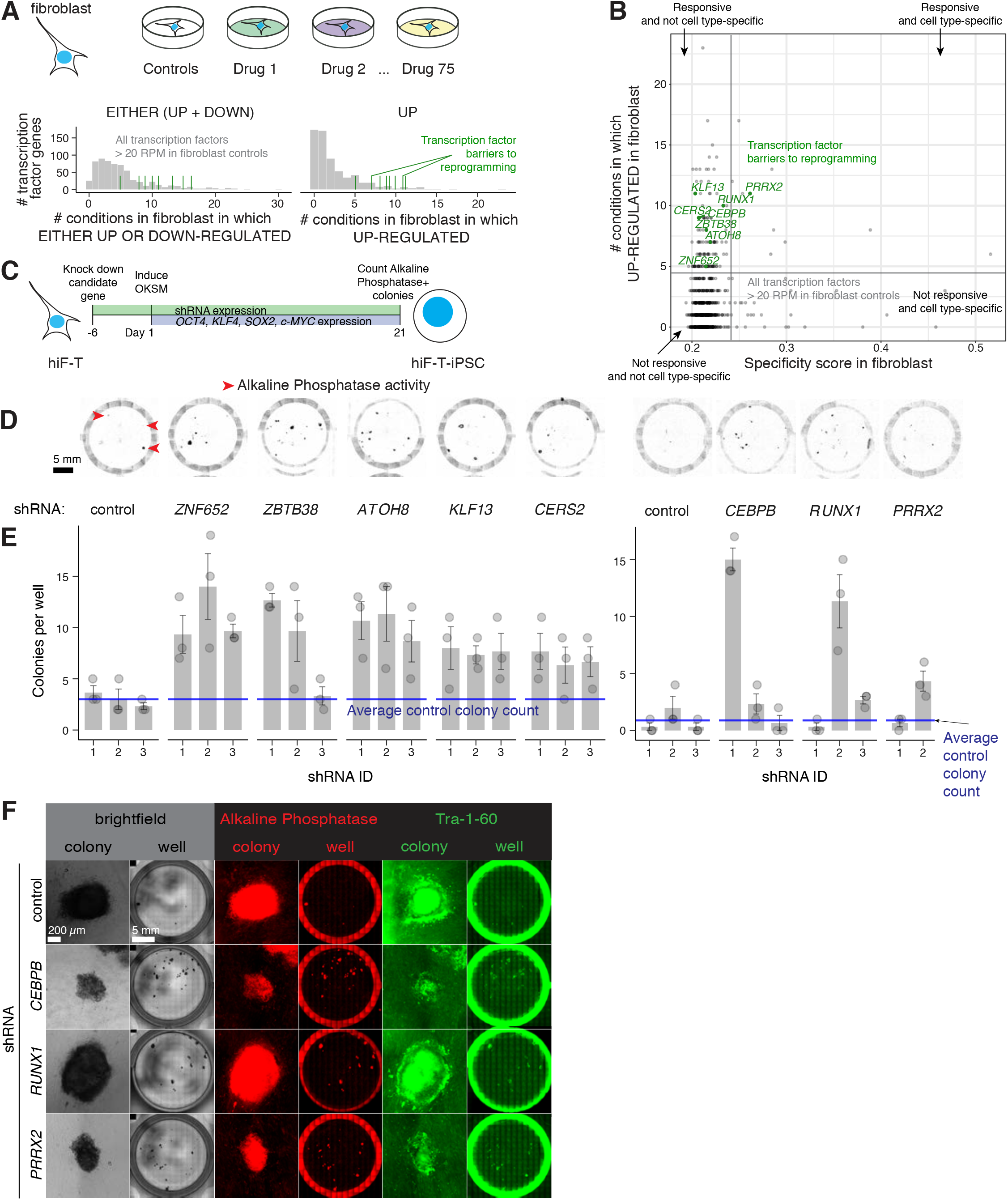
Suppression of fibroblast-responsive genes in a directed change of cell identity. A) P^3^ schematic (top) and responsiveness results (bottom) for primary human GM00942 fibroblasts (primary fibroblasts) exposed to 75 different perturbation conditions and vehicle control. All transcription factors expressed at 20 RPM or greater in vehicle control fibroblasts (grey histogram). Selected responsive genes used in later reprogramming experiments shown in green. B) Comparison of responsiveness and primary fibroblast-specificity score per gene (Jensen-Shannon specificity to primary fibroblasts without GTEx Heart and Skin, See Methods). Quadrant-separating lines chosen as elbows of the cdf of each feature over all transcription factor genes expressed at 20 RPM or greater in controls. Transcription factor barriers to reprogramming are 8 of 16 factors tested found to have a significant effect (Fisher’s exact test p-value < 0.05) on colony count against controls. See Methods for a full list of factors tested. C) Schematic of experimental workflow for hiF-T fibroblast reprogramming experiments. In parallel, hiF-Ts were transduced with shRNAs targeting genes perturbable in fibroblasts (3 shRNAs per target, each cultured in technical triplicate per experimental replicate), expanded in culture for 6 days, counted, and seeded in parallel on MEF feeders. Dox-mediated OKSM induction for 20 days prior to fixation and analysis of reprogrammed colonies. D) Representative fluorescence microscopy images of Alkaline Phosphatase activity (Vector Red staining, 4X magnification, 555 nm wavelength channel) in reprogrammed hiF-Ts under different knockdown conditions after 20 days of OKSM induction. Activity shown in black, as indicated by red arrowheads. Scale bar 5 mm. E) Colony counts with alkaline phosphatase activity after reprogramming for representative experimental replicates for two batches of perturbable target genes. F) Further characterization of hiF-T-iPSC colonies after reprogramming under different knockdown conditions. 10X magnification brightfield images (top 2 rows), Alkaline Phosphatase activity by Vector Red (middle 2 rows), and Tra-1-60 expression as identified by immunofluorescence (bottom 2 rows) shown for the same wells and inset colonies. Scale bar 5 mm for wells, 200 μm for colonies.

To test whether the set of responsive transcription factor genes was enriched for factors whose downregulation could enhance reprogramming efficiency, we first chose 16 transcription factors that were often responsive in fibroblasts in our P^3^ experiments, regardless of their cell type-specificity to fibroblasts. We considered any transcription factor genes that were 1) up-regulated in at least 5 conditions in GM00942, 2) up-regulated in at least 50% more conditions than they were down-regulated in GM00942, 3) not also frequently up-regulated in iPSC-derived cardiac myocytes, 5) expressed at >50 TPM in GM00942 controls, 6) not commonly studied in the context of fibroblast development based on literature review, 7) not commonly considered a member of a stress response or apoptosis pathway based on literature review, and 8) with at least 3 quality-controlled targeting shRNA clones available through our university core service lab’s Human TRC 2.0 lentivirus library (See Methods). Out of the set of 66 highly responsive factors identified by P^3^ in GM00942, 41 met all of the above criteria, of which we tested 16 for practical experimental scale considerations. We selected these 16 factors for validation based on them having either the most conditions in which it was up-regulated or the highest ratio of conditions in which it was up-vs. down-regulated. The human fibroblast line we used to test for reprogramming efficiency changes was the hiF-T line (*29*), which has an integrated cassette encoding the Yamanaka factors (*OCT4, SOX2, KLF4, c-MYC*; OSKM (*30*)) under doxycycline-inducible control. We used shRNAs to knock down the expression of these 16 factors one at a time in hiF-T cells prior to the induction of OSKM, then induced OSKM to check for differences in reprogrammability. (**Fig. 2C, S9**, **Table S2**). Of these 16 transcription factors, we found that 8 increased the number of Alkaline Phosphatase-positive colonies after 3 weeks of OSKM expression (**Fig. 2B,D,E; S9 and S10**). We further confirmed reprogramming by staining for Tra-1-60, a surface marker of induced pluripotent stem cells (*31*). We found that colonies positive for Alkaline Phosphatase activity also express Tra-1-60 (**Fig. 2F**). Therefore, transcription factor gene expression responsiveness in fibroblasts identified genes important in maintaining fibroblast identity.

## Discussion

The ubiquity of transcriptomic measurements has enabled us to profile the expression levels of genes across virtually all tissue types and, soon enough, all cell types. However, while these expression profiles may reveal which factors are associated with a particular cell type, such factors may or may not play a functional role in maintaining said cell type. We have found that high responsiveness of expression to perturbations may be a way to identify factors that play this functional role; i.e., these factors are “springs” that pull cells back to their identity, rather than “pillars” that remain constant in the face of perturbations.

The most prevalent way to identify candidate factors for maintaining cell type is to analyze compendiums of transcriptional profiles across various tissue types and look for genes whose expression or activity is cell type- or tissue-specific. In principle, responsiveness could have identified a proper subset of the cell type-specific genes, a superset of the cell type-specific genes, a wholly distinct set of genes, or a partially overlapping subset. We found that responsiveness seemed to identify a superset of the cell type-specific genes, meaning that there were many genes whose expression was responsive but not particularly cell type-specific. When we explicitly tested whether these responsive-but-not-cell type-specific genes could affect maintenance, we found that, for fully half the factors we tested, suppression of the factor led to an increase in reprogramming efficiency. This validation rate suggests that these responsive factors are indeed important for cell type maintenance and not merely falsely identified by P^3^. The fact that these non-cell type-specific factors are important for maintenance suggests that cell type maintenance may involve factors that act as general barriers to cell type transformations in addition to factors specific to particular types. (Although it is possible that such factors may only act as barriers in particular cell types.) Our results indicate that there are many more cell type maintenance factors than cell type-specificity alone would suggest.

It is important to note here that cellular identity maintenance may be only one of many important roles that is associated with responsive transcription factors. Our “springs” model predicts that transcription factors that maintain identity are much more likely to be responsive to perturbations. However, there are a large number of potentially overlapping functions that any given transcription factor can have, such as identity-establishment, critical cellular behaviors, and several others. Determining which of these properties most strongly associates with responsiveness would require extensive functional testing of all such properties together with P^3^.

We posit there may be at least two reasons why identity-maintaining factors may be highly responsive to perturbation. One possibility is that identity-maintaining factors are simply downstream of more signaling pathways than other transcription factors in the cell type that they maintain, hence their increased frequency of responsiveness to a panel of perturbations. Given that the responsiveness of genes was different in the two cell types that we performed P^3^ in, such a rationale would require the often invoked “cell type-specific regulatory differences”, however. Another possibility is that the upregulation of these genes is part of a feedback mechanism that mediates a homeostatic response to a perturbation. Such a response may be particular to specific signals, or may be part of a more general response to a perturbative change in some aspect of the physiology or regulation of a cell’s identity. Given that the genes identified by P^3^ were responsive to several perturbations, we suspect the latter. Another interesting question about responsiveness is whether the responses are transient or sustained. The presence of feedback loops or other homeostatic mechanisms may temper the signal over time, leading to transient responses, whereas the lack of such mechanisms may yield sustained responses. Temporal measurements may help distinguish between transient and stable responses.

It is important to distinguish between identity-establishing factors versus identity-maintaining factors. Although some identity-establishing genes are also identity-maintaining, such as *GATA4* and *TBX5* in cardiac myocytes, *OCT4* in pluripotent stem cells, and *MYT1L* in neurons (*24*, *32*, *33*), the converse need not be frequently true. That is, under the interpretation that responsiveness is a hallmark of identity maintenance, then one could assume that responsive factors would not necessarily also help establish a new cellular identity altogether. Consistent with this logic, we found that the transdifferentiation of fibroblasts to cardiac myocytes was not enhanced by adding responsive factors to the existing seven factor (7F) cocktail (**Fig. S12**). (Technically, however, it is far more difficult to evaluate the effects of overexpressing large combinations of factors (*12*, *34*–*37*)). Other approaches may be required to solve that complementary question.

An alternative approach that has been used to find cellular identity-maintaining factors is genetic or small molecule screens (*29*, *38*–*41*). Cacchiarelli et al. used pooled shRNA screens targeted towards a set of epigenetic factors to identify a number of such maintenance factors. It is difficult, however, to perform such screens in a genome-wide fashion owing to technical considerations. Other groups have screened small molecules that may affect reprogramming, and one such effort discovered the importance of *PRRX1*, a family member of *PRRX2,* the latter of which we found to maintain fibroblast identity (*41*). We envision that P^3^ may serve as a useful complement to such screens by providing a means by which one could identify on the order of a hundred candidate factors, which could subsequently be tested in a pooled screen. Such a combined approach may allow for rapid and unbiased identification of identity-maintaining factors.

## Materials and Methods

### Cell culture

Unless otherwise noted, all cell culture incubations below were performed at 37°C, 5% CO_2_. We tested intermittently for mycoplasma contamination.

#### GM00942 human dermal fibroblast culture

We cultured GM00942 human dermal fibroblasts (Coriell, GM00942; XX donor, normal-appearing tissue) according to the distributor’s instructions, on tissue culture-treated dishes in E-MEM (QBI 112-018-101) + 10% FBS (Life Technologies 16000044, lot 1802004) + Pen/Strep.

#### GM11169 human cardiac fibroblast culture

We cultured GM11169 human cardiac fibroblasts (Coriell, GM11169; XX donor) on tissue culture-treated dishes in DMEM w/Glutamax + 9% FBS (Life Technologies 16000044) + P/S.

#### HEK293FT culture

We expanded HEK293FT cells in DMEM w/Glutamax + 9% FBS + P/S.

#### hiF-T culture

We cultured hiF-T cells as previously described prior to hiF-T-iPSC reprogramming experiments(*29*). Briefly, we expanded hiF-T cells in growth medium on TC plastic dishes coated with Attachment Factor (Fisher S006100), and split cells 1:3 when they reached 60-70% confluency. hiF-T growth medium (GM) is DMEM/F-12 w/ Glutamax (Life Tech. 10565018) + 10% ES-FBS (Life Tech. 16141079) + 1x 2-Mercaptoethanol (Life Tech. 21985023) + 1x NEAA (Invitrogen 11140050) + P/S + 0.5μg/mL Puromycin + 16ng/mL rhFGF-basic (Promega G5071).

#### Immortalized human cardiac fibroblast (immHCF) culture

We cultured immortalized human cardiac fibroblasts (HCFs) as previously described (*22*). We received HCFs from Deepak Srivastava (Gladstone Institutes/UCSF). In brief, we expanded HCFs on gelatin-coated (Millipore ES-006-B) tissue culture dishes in iCM medium (per 500mL: 350mL DMEM w/Glutamax + 85mL Medium 199 + 50mL FBS + 5mL Non-essential amino acids + 10mL Pen/strep).

#### Platinum-A (Plat-A) retroviral packaging cell line culture

We cultured Platinum-A cells (Cell Biolabs RV-102) according to the manufacturer’s instructions. We expanded these cells in DMEM w/Glutamax + 9% FBS + P/S + 10ug/mL Blasticidin + 1μg/mL Puromycin.

### Cellular reprogramming

#### Derivation of cardiac myocytes (iPSC-CMs) from iPS-GM942-SeV3 iPSCs

We differentiated cardiomyocytes from iPS-GM942-SeV3 cells to create iPS-GM942-SeV3-CMs, as previously described(*42*–*46*). Throughout this manuscript we refer to iPS-GM942-SeV3-CMs as ‘iPSC-derived cardiac myocytes’ or ‘iPSC-CMs’. Briefly:

#### Seeding of iPS-GM942-SeV3

We thawed iPS-GM942-SeV3 and grew them in feeder-free conditions on Geltrex-coated (ThermoFisher, cat. A1413301) dishes in StemMACS iPS Brew-XF (Miltenyi, cat. 130-104-368) for 4-5 days, until ~75% confluency. Then, we split and seeded iPS-GM942-SeV3 into Geltrex-coated 12-well plates at a density of 3 × 10^5^ cells per well in iPS-Brew + 2μM Thiazovivin (Sigma, cat. SML1045-5MG). After 24 hours, we changed the culture medium to iPS-Brew + 1μM Chiron 99021 (Cayman Chemical, cat. 13122) and then incubated the cells for an additional 24 hours.

#### Differentiation to iPS-GM942-SeV3-CMs (a.k.a. iPSC-CMs)

Starting on Day 0, we incubated iPS-GM942-SeV3 for 18 hours in RPMI (Life Technologies, cat. 11875-119) + 100ng/ml Activin A (R&D systems, cat. 338-AC-010) + 2% B-27 (minus insulin; Life Technologies, cat. 17504-044). Next, on Day 1 we changed the medium to RPMI + 2% B-27 (minus insulin) + 5ng/ml BMP4 (Peprotech, cat. AF-120-05ET) + 1uM Chiron 99021 and incubated these cells for 48 hours. On Day 3 we changed the medium to RPMI + 2% B-27 (minus insulin) + 1uM Xav 939 (Tocris Bioscience, cat. 3748) and incubated the cells for 48 hours. On Day 5 we changed the medium to RPMI + 2% B-27 (minus insulin) and incubated the cells for 72 hours. From Day 8 - 12 we changed the medium to RPMI + 2% B-27 (including insulin) + 1% pen/strep, and replaced the medium every other day.

#### Glucose-free medium selection steps for cardiac myocytes

In order to enrich the culture for cardiac myocytes, we subjected these cells to two low glucose selection steps. On Day 12 we started the first selection step by replacing the medium with RPMI glucose free (ThermoFisher, cat. 11879020) + 2% B-27 (including insulin) + 1% Pen-Strep and incubated for 72 hours. On Day 15 we replated the cells onto Geltrex-coated dishes at 6.3e5 cells/cm^2^ in RPMI + 20% FBS (Seradigm, lot 050B14) + 1uM Thiazovivin and incubated them for 24 hours. On Day 16 we changed the medium to RPMI + 2% B-27 (including insulin) and incubated them for 48 hours to recover. On Day 18 we started the second selection step, again by replacing the medium with RPMI glucose free (ThermoFisher, cat. 11879020) + 2% B-27 (including insulin) + 1% Pen-Strep and incubated for 72 hours.

#### Replating iPS-GM942-SeV3-CMs into 96-well plates

On Day 21 of differentiation, we passaged the iPS-GM942-SeV3-CMs into Geltrex-coated 96-well plates at a density of 1e5 cells per well in RPMI + 20% FBS + 1μM thiazovivin and incubated overnight. On Day 22 we changed the medium to RPMI + 2% B-27 (with insulin) + antibiotics and cultured for an additional 48 hours to allow the plated cultures to start contracting again. By Day 23 or 24 we expected to observe recovery of contractile activity among a majority of the cardiac myocytes in all of the wells. If we did not observe beating activity we changed the RPMI + 2% B-27 (with insulin) + pen/strep medium on Day 24, incubated for another 48 hours, and checked again for contractile activity on Day 26. If the cardiac myocytes did not regain contractile activity by Day 26, we did not proceed to perturbation culture.

### Perturbation culture

#### GM00942 fibroblasts

For GM00942 fibroblasts, we called the day on which they were seeded in 96-well plates Day −2 of perturbation. We split 95% confluent 10cm tissue culture-treated dishes of cells into 96-well plates at a density of roughly ~1.5e4 cells per well in EMEM + 10% FBS + Pen/strep and incubated for 48 hours. On Day 0 we replaced the medium (250 μL) in each well and added 0.7μL of drug stock in DMSO (see **Table S1**) or of DMSO (for control cultures). This kept total DMSO concentration of the perturbation culture medium below 0.3% for all conditions. We incubated cells for 48 hours, and then on Day 2 replaced medium and re-adding a fresh dose of drug stock at the same volumes as Day 0. We incubated cells for a further 48 hours before taking images of each well (below) and extracting RNA (below) on Day 4.

#### iPSC-derived cardiac myocytes

For iPSC-CMs, we called the first day on which we observed beating activity in the majority of wells of each 96-well plate Day 0 of perturbation. On Day 0 we replaced the medium with 250μL RPMI + 2% B-27 (with insulin) + pen/strep and added 0.7μL of drug stock in DMSO (See **Table S1**) or of DMSO (for control cultures), again keeping total DMSO concentration of perturbation culture medium below 0.3% for all conditions. We incubated cells for 48 hours, and then on Day 2 replaced medium and re-adding a fresh dose of drug stock at the same volumes as Day 0. We incubated cells for a further 48 hours before taking videos of each well (below) and extracting RNA (below) on Day 4.

##### Transdifferentiation of fibroblasts to induced cardiac myocyte-like cells

We performed transdifferentiation of immortalized HCFs (immHCF), GM11169 fibroblasts, and GM00942 fibroblasts to induced cardiac myocyte-like cells as previously described(*22*). Briefly, on day −3 we plated 10^4^ fibroblasts per well in 12-well culture vessels in iCM medium and Plat-A cells in 10cm dishes (4e6 cells per dish) in DMEM w/Glutamax + 9% FBS without any antibiotics. On day −2 we transfected each dish of Plat-A cells with 10ug of one indicated pMXs expression plasmid in 500uL Optimem + 35uL Fugene HD. On day 0, we collected viral supernatants and pooled them as needed for replicate conditions, filtered them through 0.45μm filter units, and transduced fibroblasts. For transductions we used 6μg/mL polybrene, a 30min 930 x g spin, and overnight incubation at 37C. On day 1 we replaced transduction medium with iCM medium. On day 4 we replaced iCM medium with 75% iCM medium/25% Reprogramming medium (RPMI 1640 + B-27 + P/S), on day 7 with 50% iCM medium/50% Reprogramming medium, on day 11 with 25% iCM medium/75% Reprogramming medium, and on day 14 with Reprogramming medium alone. We then changed reprogramming medium daily until analysis on the indicated day per experiment, usually day 24. On day 24, we fixed cells in two formats for analysis. Some 12-well TC plastic wells were fixed in place using 3.7% formaldehyde and permeabilized at least overnight with 70% Ethanol in 4C, while others were dissociated with Accutase and transferred to Concanavalin-coated (Sigma C0412) 8-well Lab-Tek chambers. After 90-120 minutes, transferred samples were fixed in these 8-well chambers using 3.7% formaldehyde and permeabilized at least overnight with 70% ethanol in 4C. Samples in 12-well wells were processed for FISH imaging by excising them from their 12-well plate after fixation with a heated 20mm cork borer and processing them for FISH or immunofluorescence as described below.

##### Cloning of transcription factor genes and TurboGFP into pMXs

In order to drive overexpression of perturbable transcription factor genes and TurboGFP, we cloned cDNA for genes of interest into pMXs-gw (Addgene 18656; a gift from Shinya Yamanaka) using BP and LR Clonase II (Invitrogen). We amplified cDNA of targets of interest using attB-target-specific primers (**Table S3**). We used standard tools to verify sequence identity of the plasmid backbone and gene insert, such as restriction digestion and Sanger sequencing. We amplified attB-TurboGFP off of the SHC003 plasmid (Sigma SHC003).

##### Titering of pMXs retroviral vectors

Since our expression vectors do not contain selectable or fluorescent markers and pMXs retroviral vectors only transduce dividing cells, we indirectly titered each experimental replicate’s batch of virus by co-transducing parallel samples of HCFs with pMXs-DsRed Express (Addgene 22724; a gift from Shinya Yamanaka) and pMXs-TurboGFP (see “Cloning” above) produced using the same batch of Plat-A cells under the same conditions. In order to estimate the fraction of cells that are infected at least once per factor, we considered the infection rates of these fluorescent pMXs vectors. By comparing the fraction that are co-infected with both against the fraction that are infected with each factor at all, we can infer the fraction of cells dividing in the population during the transduction period and the fraction of those cells that receive at least one copy of any individual expression vector. We make the simplifying assumption that among dividing cells infection events are independent of each other. Therefore, the ratio of the fraction that are DsRed+ and GFP+ to the fraction that are DsRed+ (or GFP+) is approximately the square of the transduction rate for any individual virus. E.g., for 30% of cells being DsRed+ and 24.3% being DsRed+ and GFP+, 24.3/30 = 0.81, which gives 90% transduction rate for each individual virus. We used two-color flow cytometry (GUAVA) to assess DsRed+, GFP+, and DsRed+ GFP+ fractions per sample.

##### shRNA-mediated knockdown of transcription factors in hiF-T cells

We conducted knockdown of individual transcription factors using shRNAs essentially as previously described.(*29*) In brief, we acquired cloned pLKO.1, pLKO.1-shRNA, and pLKO.1-TurboGFP plasmids from the University of Pennsylvania High-Throughput Screening Core (**Table S2**). We verified shRNA and backbone sequence with Sanger sequencing. We packaged shRNA lentivirus using pMD2.G (Addgene 12259; a gift from Didier Trono) and psPAX2 (Addgene 12260; a gift from Didier Trono) in HEK293FT cells, and filtered viral supernatant through 0.22μm filter units prior to infecting hiF-T cells. We infected hiF-T cells at an MOI of approximately 1 (for a transduction efficiency of ~70%) with 4μg/mL polybrene and 30 min 930 x g centrifugation. Since hiF-T cells are already Puromycin-resistant, we were unable to perform an antibiotic selection step after infection with these pLKO.1-puro-based shRNA plasmids.

#### Verification of knockdown efficiency following shRNA transduction

We performed RT-qPCR on RNA extracted from samples of the hiF-T cells that we used in reprogramming experiments. We used Superscript III Reverse Transcriptase for first-strand cDNA synthesis and Power SYBR qPCR Master Mix with gene-specific primer pairs for qPCR on an Applied Biosystems 7300 system. We performed all statistical analysis using custom scripts in R (see “Code accessibility” below for all scripts) and calculated knockdown efficiency using the ΔΔCt method.

##### hiF-T reprogramming to pluripotency

We performed hiF-T reprogramming experiments as previously described. Briefly, after shRNA transduction on day −7, we expanded cells in hiF-T GM without puromycin for one week. On Day −1 we seeded CF-1 Irradiated MEFs on uncoated 24-well plates (Corning) at a density of 2.5 × 10^5^ cells per well in hiF-T GM without puro. On Day 0, we seeded 10^4^ hiF-T cells per 24-well plate well. On Day 1 we began Yamanaka factor induction by switching media to hiF-T GM with 2μg/mL doxycycline and without puromycin. On Day 3 we switched media to KSR Medium (KSRM): DMEM/F-12 w/ Glutamax (Life Tech. 10565018) + 20% Knockout Serum Replacement (Life Tech. 10828010) + 1x 2-Mercaptoethanol (Life Tech. 21985023) + 1x NEAA (Invitrogen 11140050) + P/S + 8ng/mL rhFGF-basic + 2μg/mL Doxycyclin. We changed KSRM daily, and analyzed cells on day 21. We performed a total of 3 biological replicates (i.e., different vials of hiF-T cells expanded and reprogrammed on different days with different batches of media), each with technical triplicates per shRNA condition. All biological and technical replicates of all reprogramming experiments we conducted are presented in this manuscript.

##### High-throughput RNA extraction

We used RNaqueous-96 kits (Ambion AM1920) for RNA extraction without the optional DNase step, according to manufacturer’s instructions.

##### RNAtag sequencing

We conducted highly parallelized bulk RNA sequencing with RNAtag-seq as previously described, using all components and steps in the published protocol (*20*). We ordered the specified 32 barcoded DNA oligos for RNAtags from Biosearch Technologies and indexed primers for library amplification and reverse transcription from IDT. We sequenced all RNAtag-seq libraries in batches of 96 samples on an Illumina NextSeq 550 using 75 cycle high-output kits (Illumina 20024906).

##### RNAtag-seq data processing

We demultiplexed RNAtag-seq reads using custom scripts, courtesy of Edward Wallace (*47*). We aligned RNAtag-seq reads to the human genome (hg19) with STAR v2.5.2a and counted uniquely mapping reads with HTSeq v0.6.1 (*48*). We performed all downstream analysis in R v3.6.1 using packages yaml_2.2.0, DESeq2_1.24.0, SummarizedExperiment_1.14.0, DelayedArray_0.10.0, BiocParallel_1.18.0, matrixStats_0.54.0, Biobase_2.44.0, GenomicRanges_1.36.0, GenomeInfoDb_1.20.0, IRanges_2.18.1, S4Vectors_0.22.0, BiocGenerics_0.30.0, e1071_1.7-2, magrittr_1.5, ggrepel_0.8.1, ggplot2_3.2.0, tibble_2.1.3, tidyr_0.8.3, and dplyr_0.8.3, clusterProfiler_3.12.0, org.Hs.eg.db_3.8.2, and their associated dependencies.

##### Gene expression perturbation-responsiveness

As a measure of gene expression perturbation-responsiveness we used the count of the number of conditions in which a gene was differentially expressed relative to cell type DMSO controls. For most analyses we used any change with a DESeq2 adjusted p-value less than 0.1, but also conducted analyses with additional filters, such as minimum absolute values of log2FoldChange. Filter criteria are explicitly noted in figure legends for each analysis. We also considered other measures of perturbation-responsiveness, as well, which are not explored in the manuscript above, details of which are available upon request.

##### Gene expression cell type-specificity analysis

We calculated Jensen-Shannon divergence-based scores for each gene as previously described (*28*) in each tissue or cell type in the GTEx V7 RNA-seq dataset (*27*). Briefly, we analyzed all V7 GTEx RNA-seq samples with: RIN (RNA integrity score) ≥ 7, mapping rate ≥ 50%, and 1 million or more mapped reads. With these filters, 3623 samples remained in the processed GTEx dataset. We then calculated JS_sp_ values for each gene in each tissue type (SMTS identifier) based on average expression levels per tissue type in transcripts per million (TPM), as described in Cabili et al., 2011. For the analysis presented in main text figures, we removed GTEx Skin and Heart from the dataset and added GM00942 fibroblast and iPSC-CM cardiac myocyte control average expression expression values. We present specificity score calculations with and without removal of GTEx Skin and Heart and with and without addition of GM00942 fibroblast and iPSC-CM cardiac myocyte data in the supplement.

##### Gene set enrichment analysis

We performed gene set over-enrichment analysis on highly expressed genes (mean RPM > 20) for each cell type using the clusterProfiler v3.12.0 package in R v3.6.1 (*49*). We compared iPSC-CM-responsive highly expressed genes (up-regulated in 4 or more conditions and mean RPM > 20 in control iPSC-CM) against the entire set of genes expressed > 20 RPM in iPSC-CM controls. We compared iPSC-CM-unresponsive genes (up-regulated in 0 or 1 conditions and mean RPM > 20 in control iPSC-CM) against the entire set of genes expressed > 20 RPM in iPSC-CM controls. We compared fibroblast-responsive highly expressed genes (up-regulated in 4 or more conditions and mean RPM > 20 in control GM00942 fibroblasts) against the entire set of genes expressed > 20 RPM in GM00942 controls. We compared fibroblast-unresponsive genes (up-regulated in 0 or 1 conditions and mean RPM > 20 in control GM00942) against the entire set of genes expressed > 20 RPM in GM00942 controls.

##### Prioritization of highly responsive genes for use in reprogramming experiments

For iPSC reprogramming experiments we considered transcription factor genes (*50*) that were 1) up-regulated in at least 5 conditions in GM00942, 2) up-regulated in at least 50% more conditions than they were down-regulated in GM00942, 3) not also frequently up-regulated in iPSC-CM, 5) expressed at >50 TPM in GM00942 controls, 6) not commonly studied in the context of fibroblast development based on literature review, 7) not commonly considered a member of a stress response or apoptosis pathway based on literature review, and 8) with at least 3 quality-controlled targeting shRNA clones available through our university core service lab’s Human TRC 2.0 lentivirus library. Ultimately we tested knockdown of the genes ‘SKIL’, ‘YBX3’, ‘TSC22D1’, ‘CERS2’, ‘KLF13’, ‘TBX3’, ‘ID1’, ‘ATOH8’, ‘ZNF652’, ‘NFATC4’, ‘ZBTB38’, ‘LARP1’, ‘CEBPB’, ‘ID3’, ‘PRRX2’, and ‘RUNX1’.

For cardiac transdifferentiation experiments we considered transcription factor genes (*50*) that were 1) up-regulated in at least 4 conditions in iPSC-CM, 2) up-regulated in at least 50% more conditions than they were down-regulated in iPSC-CM, 3) not also frequently up-regulated in GM00942, 5) expressed at >50 TPM in iPSC-CM controls, 6) not commonly studied in the context of fibroblast development based on literature review, and 7) not commonly considered a member of a stress response or apoptosis pathway based on literature review. Ultimately we tested overexpression of the genes ‘SP3’, ‘ZBTB10’, ‘ZBTB44’, ‘SSH2’, ‘NFIA’, ‘ZNF652’, and ‘ZFP91’.

##### Live cell Tra-1-60 imaging

In a pilot reprogramming experiment without shRNA transduction, we conducted live-cell staining of hiF-T-iPSC colonies Tra-1-60 with TRA-1-60 Alexa Fluor™ 488 Conjugate Kit for Live Cell Imaging (Life Tech. A25618) according to the manufacturer’s instructions.

##### Alkaline phosphatase staining with colorimetry

We used the Vector Red Substrate kit (Vector Labs SK-5100) to stain hiF-T-iPSC colonies after fixation on day 21 of reprogramming experiments. We fixed wells in 24-well format using 3.7% formaldehyde for 3 minutes, and followed the manufacturer’s instructions.

##### Immunofluorescence

We performed immunofluorescence for several markers. For Tra-1-60 immunofluorescence of hiF-T-iPSC samples that had already been stained with Vector Red, we blocked and permeabilized in 5% BSA + 0.1% Triton X-100 in PBS at room temperature for 30 min. Then we washed samples in PBS and used Stemgent 09-0068 at 1:200 in 5% BSA + 0.1% Triton X-100 for 2 hours at room temp. We washed samples in PBS and stained with DAPI prior to imaging. For cardiac troponin immunofluorescence of iPSC-CM and transdifferentiated samples, we fixed samples in 3.7% formaldehyde for 10 min at room temp, washed in PBS, and permeabilized with 70% ethanol overnight at 4C. Independent of smFISH or after the smFISH protocol, we performed immunofluorescence with Abcam ab45932 primary (1:200) with goat anti-rabbit-Alexa 594 (1:200) secondary and Fisher MA5-12960 primary (1:200) with donkey anti-mouse-Alexa 488 (1:200) secondary. We used samples in 3% BSA + 0.1% Tween 20 for blocking/binding buffer. Primary antibody incubations of 1 hour and secondary incubations of 30 min, both at room temperature. Samples were washed with PBS and stained with DAPI prior to imaging.

##### Single-molecule RNA FISH

*GAPDH* probes were used as previously described (*51*). We incubated our cells overnight at 37°C in hybridization buffer (10% dextran sulfate, 2× SSC, 10% formamide) with standard concentrations of RNA FISH probes (**Table S3**) (*21*). The following morning, we performed two washes in wash buffer (2X SSC, 10% formamide), each consisting of a 30-min incubation at 37°C. After the second wash, we rinsed once with 2X SCC/DAPI and mounted the sample for imaging in and 2X SSC (*21*). We performed RNA FISH on cell culture samples grown on a Lab-Tek chambered coverglass using 50 μL of hybridization solution spread into a thin layer with a coverslip and placed in a parafilm-covered culture dish with a moistened Kimwipe to prevent excessive evaporation.

##### Imaging

We imaged each sample on a Nikon Ti-E inverted fluorescence microscope using a 60X Plan-Apo objective and a Hamamatsu ORCA Flash 4.0 camera. For 60X imaging of complete cells, we acquired z-stacks (0.3 μm spacing between slices). For 60X imaging of a large field of cells with one plane each, we used Nikon Elements tiled image acquisition with perfect focus. All images of stained cells were in different fluorescence channels using filter sets for DAPI, Atto 488, Cy3, Alexa 594, and Atto 647N. The filter sets we used were 31000v2 (Chroma), 41028 (Chroma), SP102v1 (Chroma),17 SP104v2 (Chroma) and SP105 (Chroma) for DAPI, Atto 488, Cy3, Atto 647N/Cy5 and Atto 700, respectively. A custom filter set was used for Alexa 594/CalFluor610 (Omega). We tuned the exposure times depending on the dyes used: 400 ms for probes in Cy3 and Alexa 594, 500 ms seconds for each probe in Atto 647N, and 50 ms for DAPI probes. We also acquired images in the Atto 488 channel with a 400 ms exposure as a marker of autofluorescence.

##### Image Processing

smFISH analysis of image scans and stacks was done as previously described using rajlabimagetools changeset 775fd10 (https://bitbucket.org/arjunrajlaboratory/rajlabimagetools/wiki/Home) in MATLAB v2019a, and is compatible with rajlabimagetools changeset a2c6ac5 (https://github.com/arjunrajlaboratory/rajlabimagetools/) in MATLAB v2017a (*21*).

##### Reproducible analysis

For all raw imaging and qPCR data, please see: https://www.dropbox.com/sh/2ny7k6c4zy6zdsh/AADLNoom3YOw0Ps0B8w509R7a?dl=0. For all raw sequencing data, please see: https://www.dropbox.com/sh/ss7eovrclone69a/AADAkowTjNZ2_FwxZSXey9rCa?dl=0. For a fully reproducible processing pipeline, including raw qPCR data, from all raw data types to the graphs and images included in this paper, please see: https://www.dropbox.com/sh/3iqo4mjx4le0a8c/AACz1J2J6Vwuq_3mLNInkYU-a?dl=0. For details, please see the readme files included in each folder.

## Acknowledgements

We thank members of the Raj lab, Jain lab, and Penn iPSC Core, particularly Eduardo Torre, Allison Cote, Aman Kaur, Ben Emert, and Kate Alexander, as well as the late John Gearhart for insightful discussions related to this work. We thank Deepak Srivastava and members of his lab, including Palmer Yu, for providing 7F vectors, immHCF cells, and advice on cardiac transdifferentiation protocols. We thank Davide Cacchiarelli and Alex Meissner for providing hiF-T cells and for advice on reprogramming protocols. We thank Mitchell Guttman, Sofia Quinodoz, and Mario Blanco for advice on RNAtag-seq. We thank Edward Wallace for making PyRNAtagSeq scripts publicly available. We thank Russ Carstens for Platinum-A cells. We thank Junwei Shi for assistance with flow cytometry. We thank the Penn iPSC Core for assistance with iPSC-derived cardiac myocytes, the Penn High-Throughput Screening Core for assistance with small molecule drugs, shRNAs, and cDNA clones, and the Penn DNA Sequencing Core for assistance with large-scale plasmid preps. The Genotype-Tissue Expression (GTEx) Project was supported by the Common Fund of the Office of the Director of the National Institutes of Health, and by NCI, NHGRI, NHLBI, NIDA, NIMH, and NINDS. The data used for the analyses described in this manuscript were obtained from the GTEx Portal on 10/05/17. The authors acknowledge grant support for this project from: NIH/NINDS F30NS100595 and NIH/NHGRI T32HG000046 to IAM, HFSP LT000919/2016-L to OS, grant from Schmidt Science Fellows (in partnership with the Rhodes Trust) to YG. YG is a fellow of the Jane Coffin Childs Memorial Fund for Medical Research and this investigation has been aided by a grant from the Jane Coffin Childs Memorial Fund for Medical Research, NIH F31HL147416 to RALS, a Penn Epigenetics Pilot Grant to PPS, WY is supported by the Penn Institute for Regenerative Medicine. Burroughs Wellcome Career Award for Medical Scientists, American Heart Association, Allen Foundation and NIH R01GM137425 to RJ, R01 CA238237, NIH Director’s Transformative Research Award R01 GM137425, R01 CA232256, NSF CAREER 1350601, P30 CA016520, SPORE P50 CA174523, NIH U01 CA227550, NIH 4DN U01 HL129998, NIH Center for Photogenomics (RM1 HG007743), and the Tara Miller Foundation to AR.

## Author Contributions

AR and IAM conceived of P^3^, and IAM, RJ, and AR conceived of the larger project with applications to cardiac transdifferentiation and reprogramming; IAM designed, conducted, and analyzed all experiments with supervision by AR and RJ, with the exception of iPSC-CM differentiation performed by HIE, RT, and WY. OS contributed to the design of some transdifferentiation experiments. WY and RT contributed to analysis of reprogramming experiments. YG contributed to some transdifferentiation experiments, analysis, and figure design. MCD, PPS, and RALS contributed to some transdifferentiation experiments. IAM designed all analyses and wrote all software, with some assistance from LEB, with supervision by AR and RJ. IAM, RJ, and AR wrote the paper.

## Supplementary figure legends

**Supplementary Figure 1:**
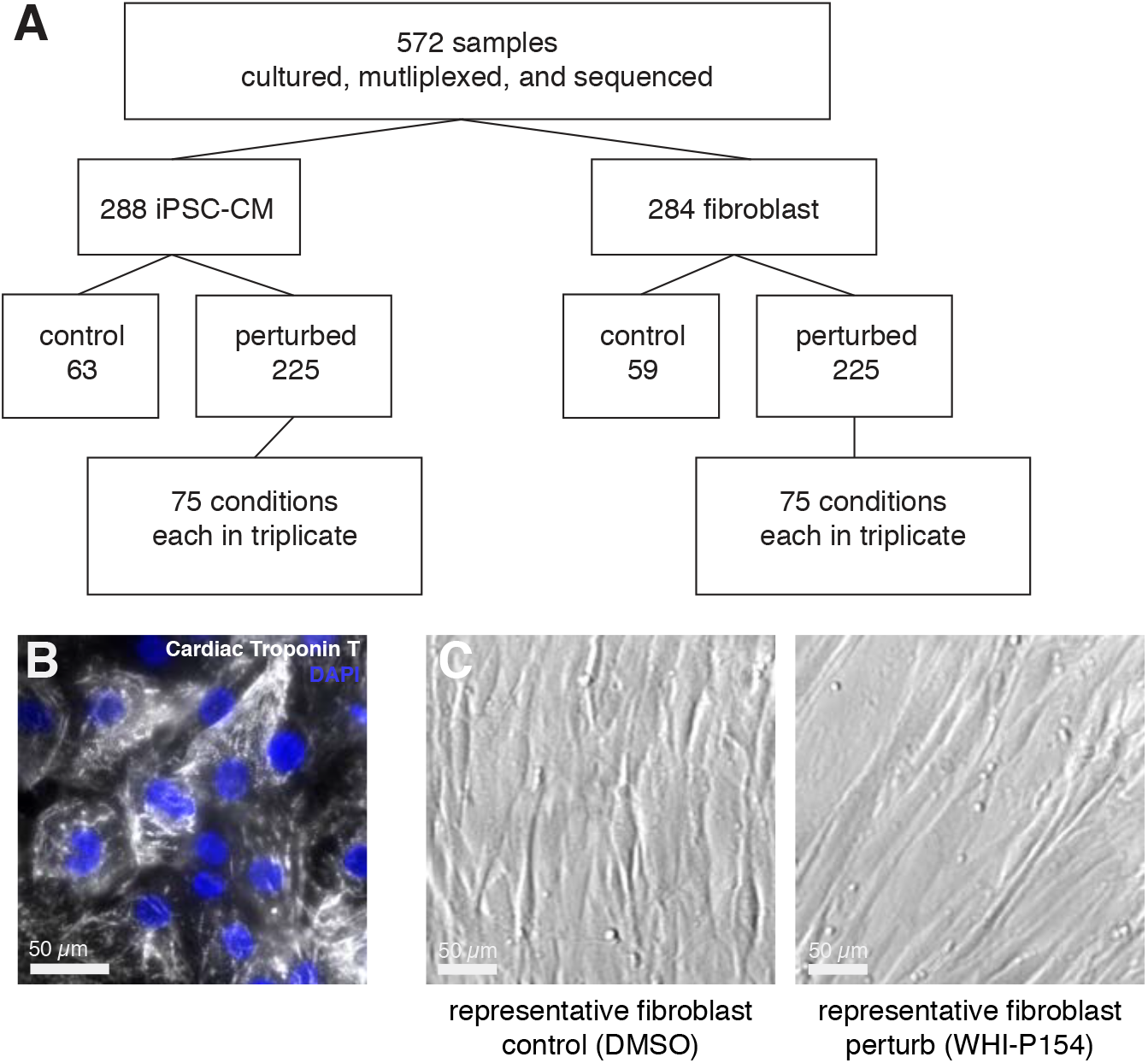
iPSC-derived cardiac myocyte and primary fibroblast samples. A) Overview of primary GM00942 fibroblast and iPSC-derived cardiac myocyte (iPSC-CM) samples processed for P3. B) Representative cardiac Troponin T immunofluorescence of iPSC-CMs (60X magnification) prior to perturbation. See Supplementary videos 1 and 2 for representative videos of control and perturbed iPSC-CM cells demonstrating contractile activity immediately prior to RNA extraction. C) Representative brightfield images (10X magnification) of control (left) and perturbed (right) primary GM00942 fibroblasts, in this image after 4 days of exposure to vehicle DMSO or WHI-P154, respectively. Scale bars 50 μm.

**Supplementary Figure 2:**
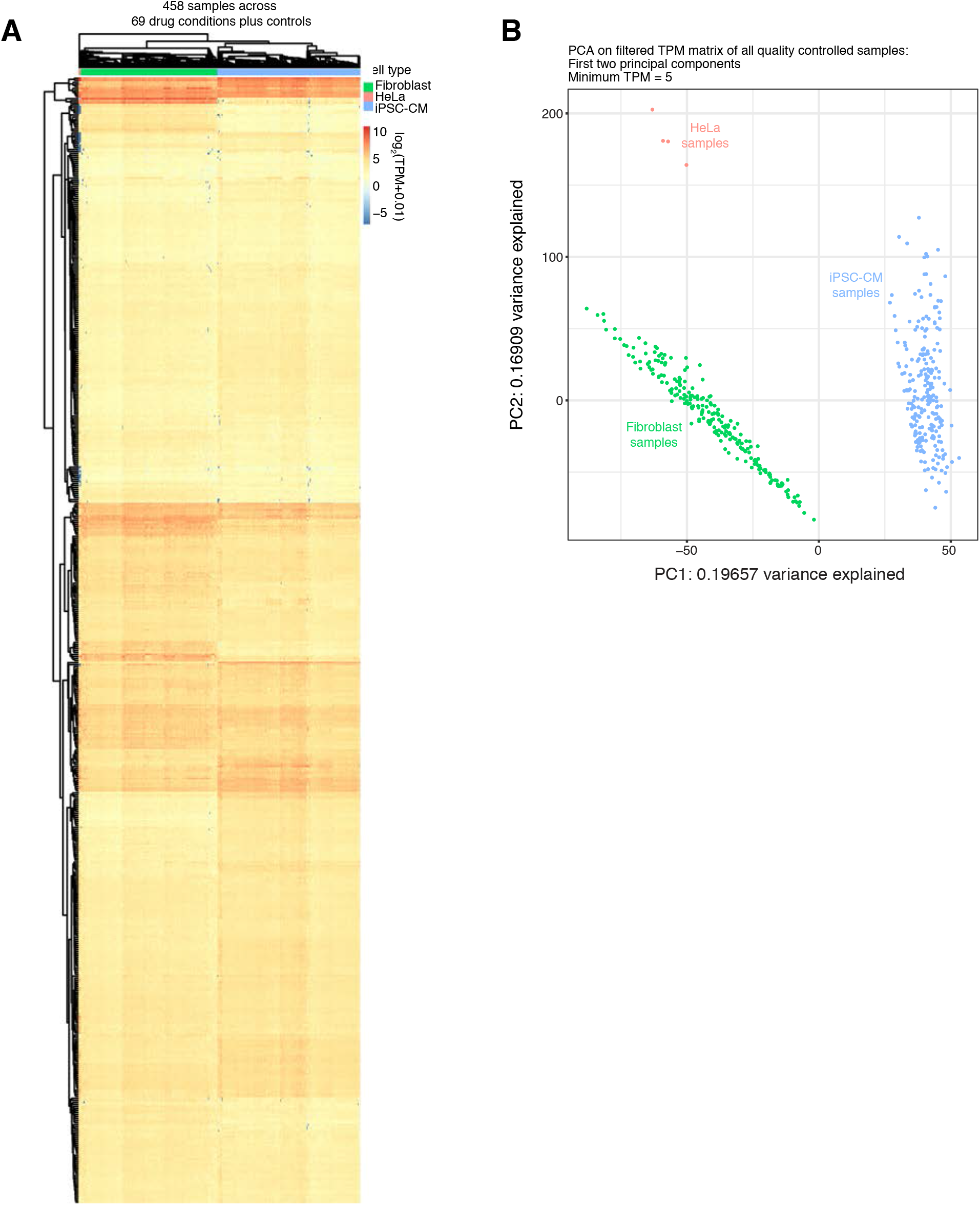
Exploratory analysis of RNAtag-seq results of perturbed iPSC-derived cardiac myocytes, primary fibroblasts, and HeLa cells. A) Heatmap of log_2_(TPM+0.01) expression values for all transcription factor genes expressed at 5 TPM or greater on average across all quality-controlled samples. B) Principal Component Analysis of all genes expressed at 5 TPM or greater on average across all quality-controlled samples.

**Supplementary Figure 3:**
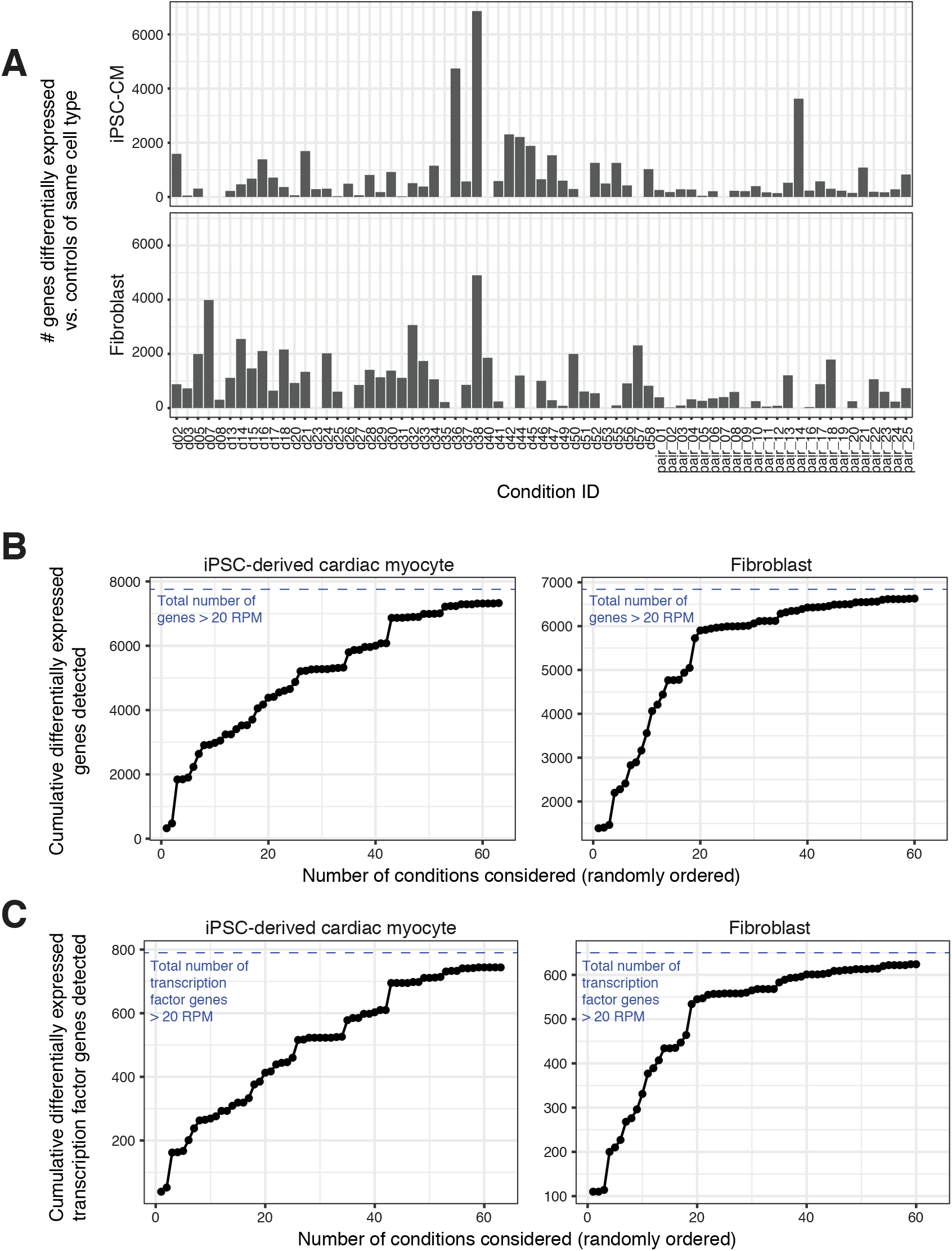
Differential expression of perturbed samples vs. controls. A) Number of differentially expressed genes (see Methods) in each condition in each cell type vs. control samples of that cell type. B) Cumulative fraction of highly-expressed genes (control RPM > 20) detected as differentially expressed in at least one condition relative to controls in the same cell type (considering all conditions, 96.9% of filtered genes differentially expressed in fibroblast and 94.5% in iPSC-derived cardiac myocytes). Blue dashed lines correspond to the number of genes expressed > 20 RPM in controls of that cell type. C) Cumulative fraction of highly-expressed transcription factor genes (control RPM > 20) detected as differentially expressed in at least one condition relative to controls in the same cell type. Blue dashed lines correspond to the number of transcription factor genes expressed > 20 RPM in controls of that cell type.

**Supplementary Figure 4:**
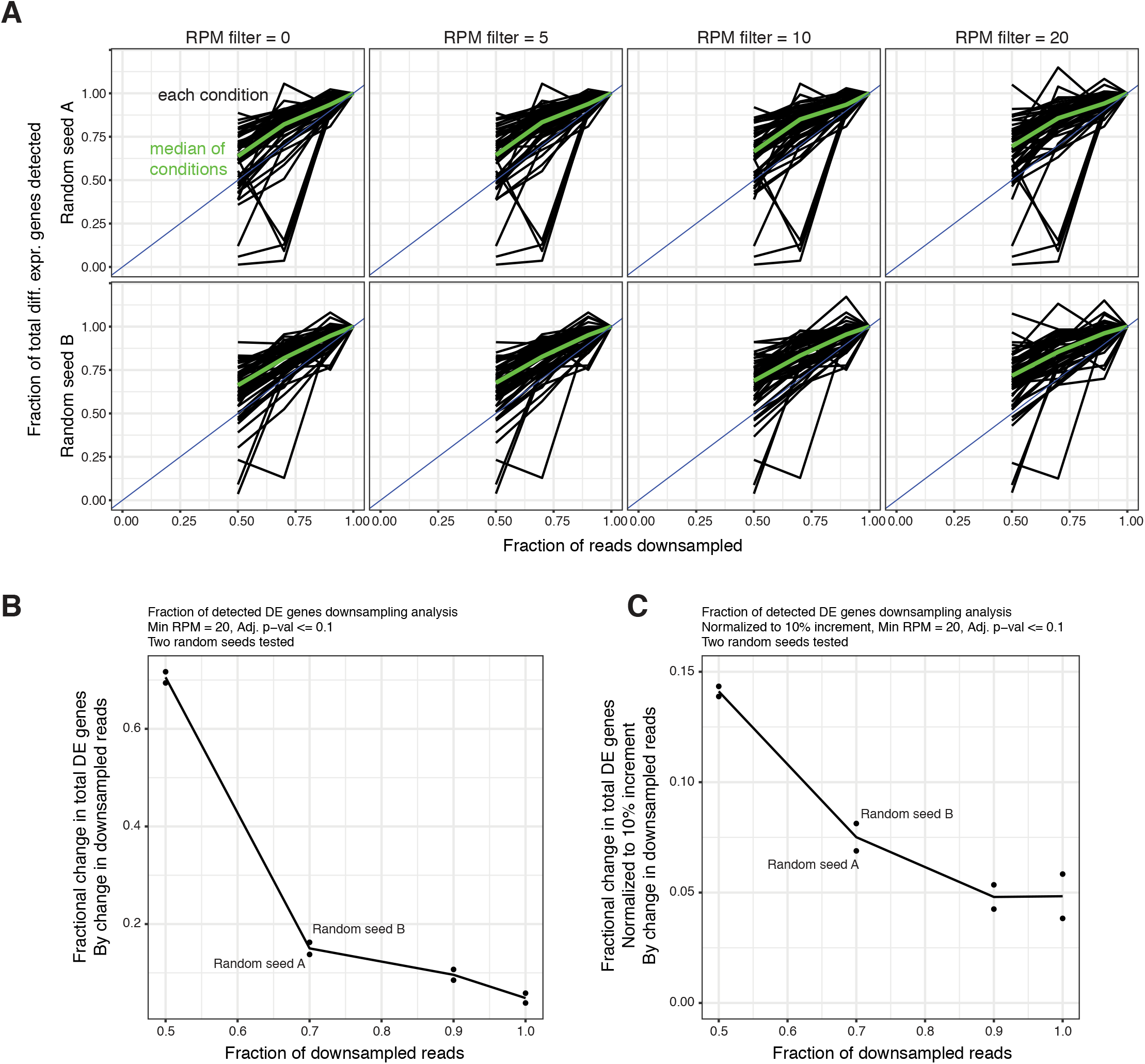
Downsampling analysis of iPSC-derived cardiac myocyte RNAtag-seq results for analyzing differential expression detection power limitations. A) Counts per sample were randomly downsampled to 50%, 70%, and 90% of the total dataset, using two different random seeds, and re-calculated. Random seed A = 2380, B = 53250. Each black line represents the fraction of differentially expressed genes still detected at the downsampled read fraction. The green line represents the median fraction of total differentially expressed genes per drug at that downsampled read fraction. Results are shown for four minimum control-average expression level filters (RPM > 0, 5, 10, and 20). Drugs are only included in this analysis if they have 200 or more differentially expressed genes detected in the full dataset. The blue line represents equal fractions of downsampled reads and detected differentially expressed genes. Some conditions show fewer differentially expressed genes after some incremental increases in sequencing depth (negative slopes) due to noise associated with low total read counts for some genes. B) Median gains in differentially expressed genes per condition detected with each simulated increment of additional reads (two simulations per fraction), as a fraction of the actual number of differentially expressed genes per condition. C) Normalized median gains in differentially expressed genes per condition detected with each simulated increment of additional reads (two simulations per fraction), estimated percent increase in number of detected differentially expressed genes per simulated additional 10% of total reads.

**Supplementary Figure 5:**
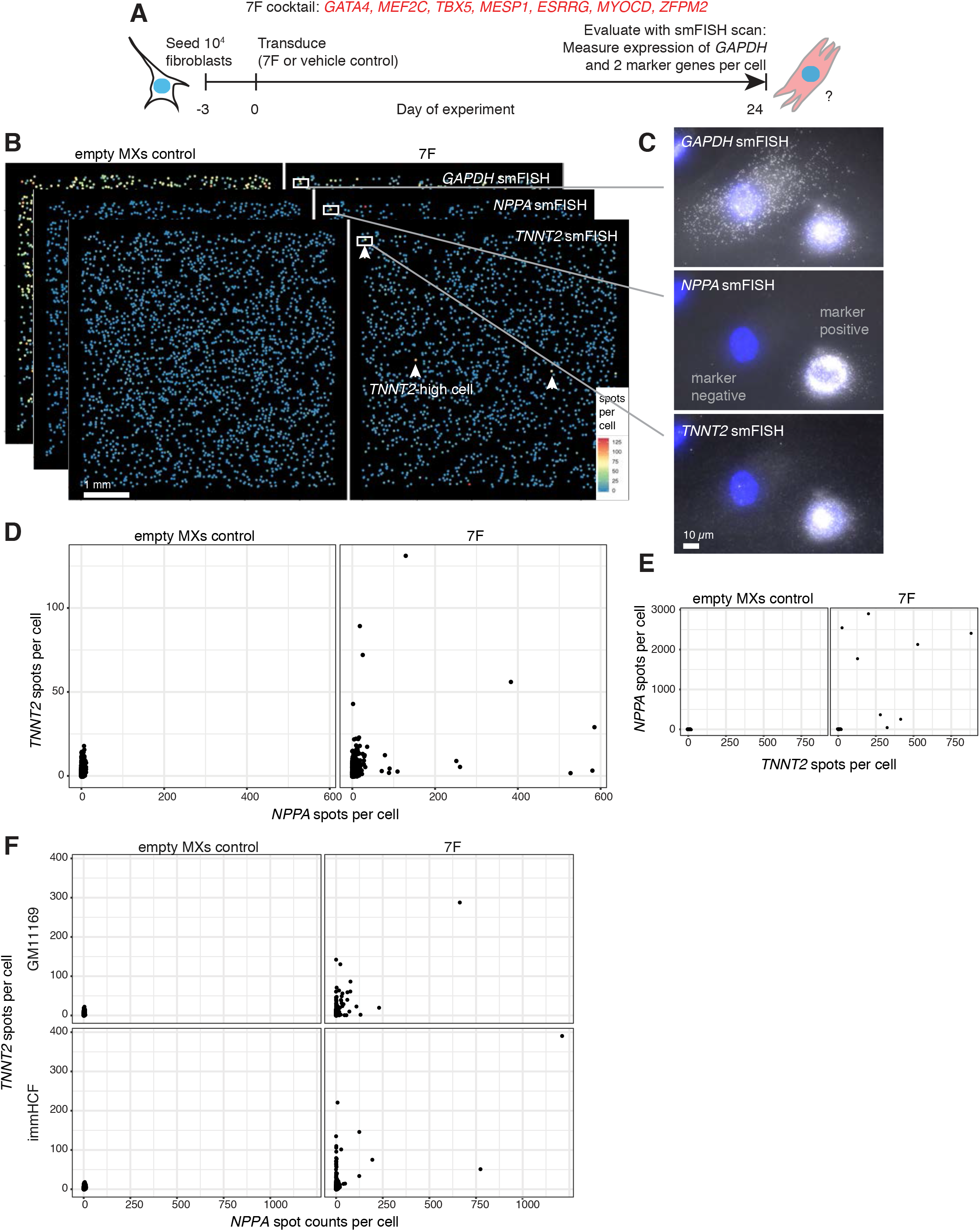
Transdifferentiation of fibroblasts to cardiac myocyte-like cells. A) Schematic of experimental workflow. See Methods for full details. Also appears in Figure 1. B) Reconstruction and per-gene quantification of smFISH scans of GM00942 cells exposed to vehicle control (MXs-empty; 2267 cells) or 7F (1693 cells). One z-slice per cell. Examples of cell with high expression of TNNT2 marked with white arrowheads. Scale bar 1 mm. C) Representative images of GAPDH, NPPA, and TNNT2 smFISH demonstrating a cell that expresses marker genes NPPA and TNNT2 at high levels and one cell that does not. Scale bar 10 μm. Also appears in Figure 1. D) Comparison of single-cell TNNT2 and NPPA expression levels for all cells in smFISH scans. E) smFISH counts per cell of two marker genes for cells after receiving either vehicle control (MXs-empty) or 7F. F) smFISH assessment of transdifferentiation of immHCF and GM11169 human cardiac fibroblasts after exposure to vehicle control (MXs-empty) or 7F. Average of 1151 +/− 51 cells per smFISH scan per condition.

**Supplementary Figure 6:**
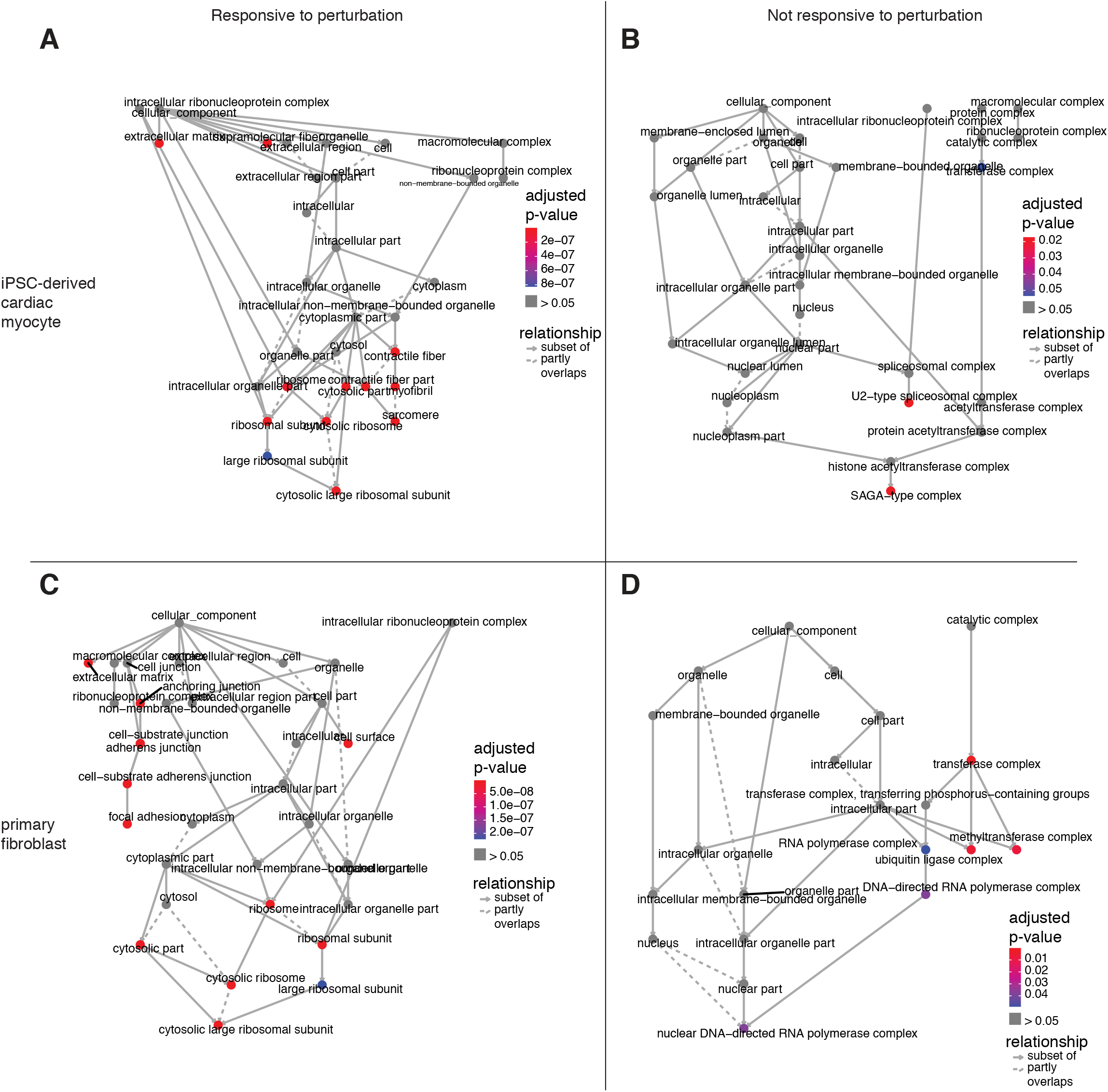
Gene set enrichment analysis of perturbed iPSC-derived cardiac myocyte and primary fibroblast samples. Gene set over-enrichment analysis was performed for all Gene Ontology - Cellular Component terms. For each cell type (A,B: iPSC-derived cardiac myocyte; C,D: primary fibroblast) genes were filtered first to all genes expressed at 20 RPM or greater in controls. GO terms were tested for over-enrichment in the set of filtered genes that are frequently up-regulated (up in 4 or more conditions, A,C) relative to all filtered genes, as well as in the set of filtered genes that are infrequently up- or down-regulated (up or down in 1 or 0 conditions total, B,D) relative to all filtered genes.

**Supplementary Figure 7:**
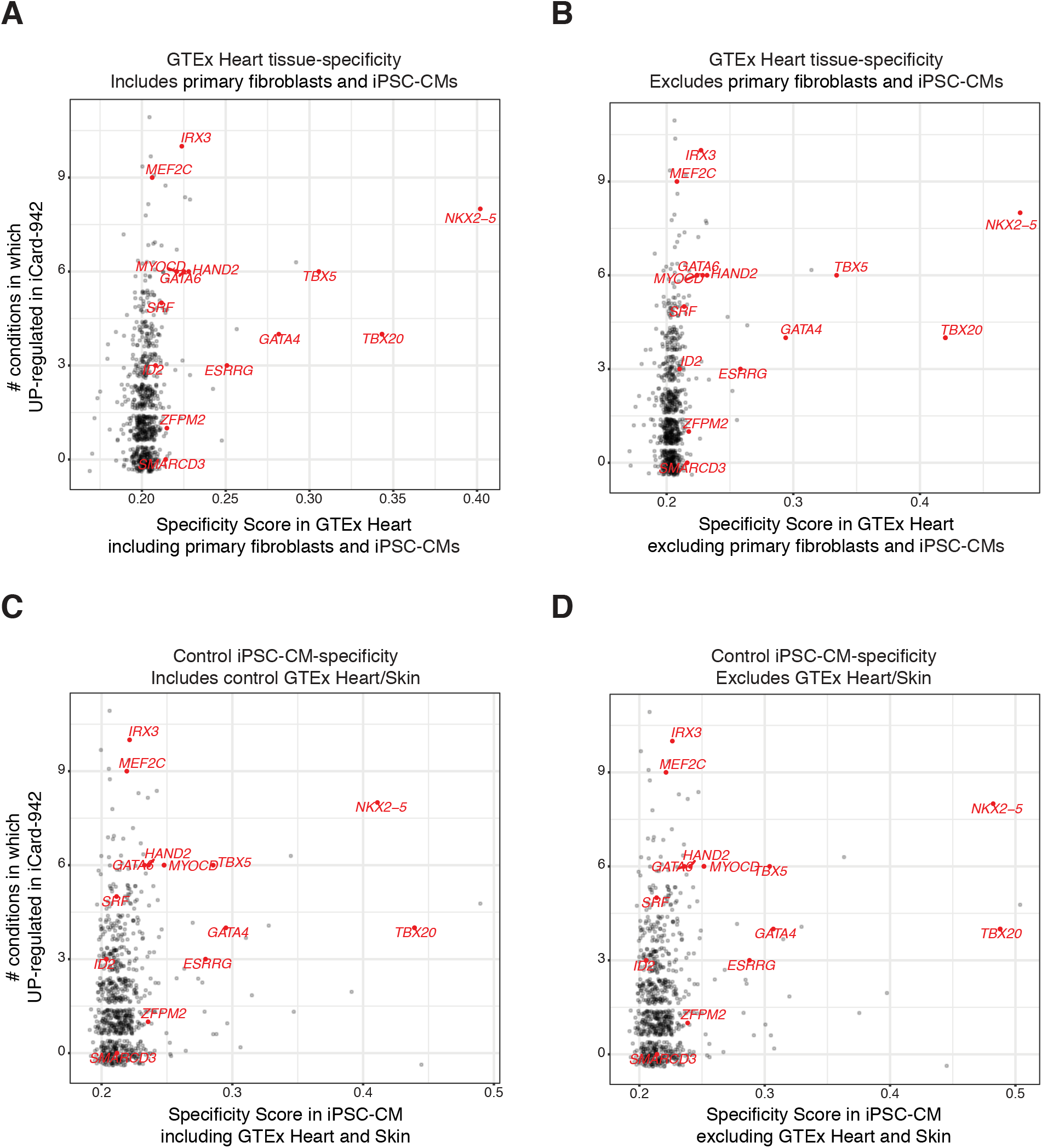
Transcription factor gene expression specificity in iPSC-derived cardiac myocytes. Expression specificity scores for heart (A,B) and iPSC-derived cardiac myocytes (C,D) are plotted on the x axis. Analyses were repeated on datasets including (A,C) or excluding (B,D) the most similar samples. Responsiveness to perturbation, i.e., the number of conditions in which a gene was expressed higher than in controls, is on the y axis. Panel D also appears in Figure 1.

**Supplementary Figure 8:**
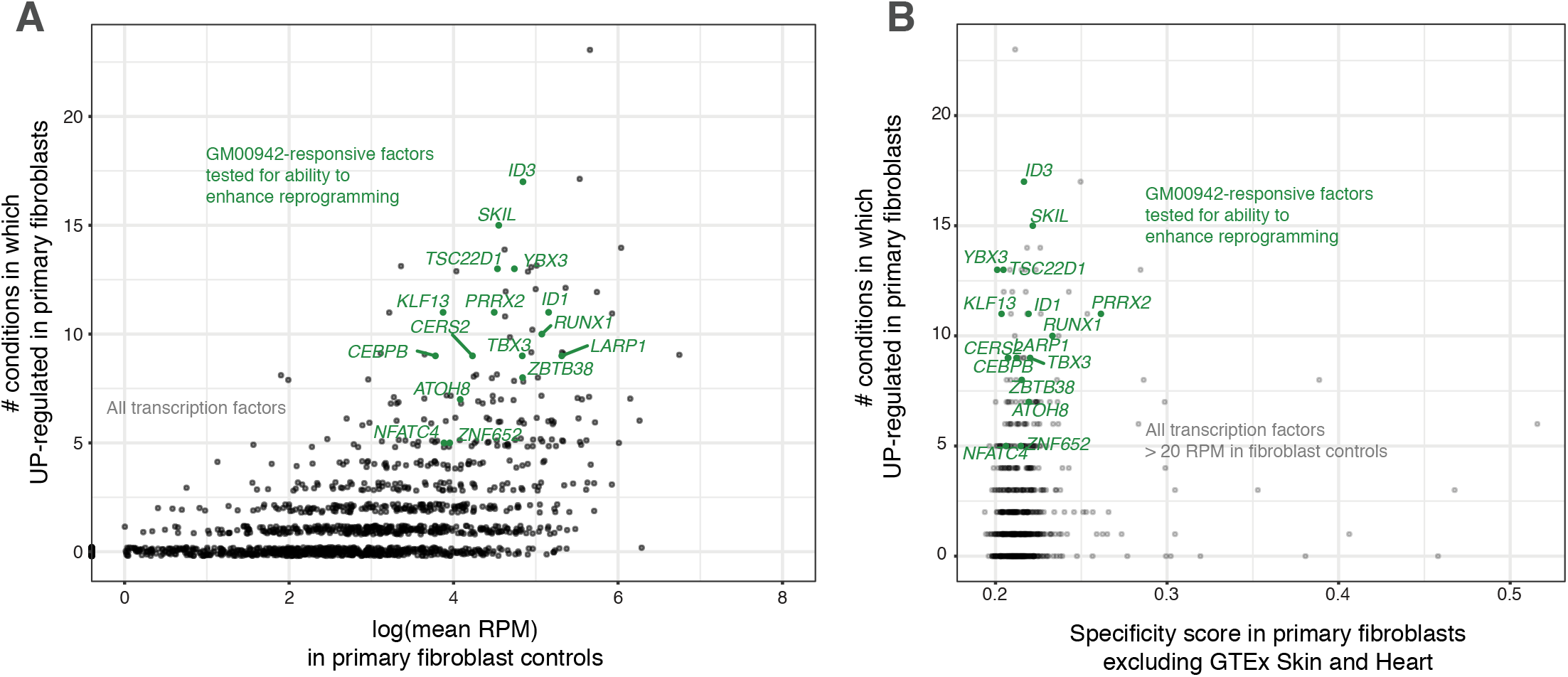
P^3^ analysis of perturbed primary fibroblasts. A) Responsiveness to perturbation (i.e., number of conditions in which up-regulated in GM00942 primary fibroblasts) against average expression in control GM00942 primary fibroblasts. B) Responsiveness to perturbation against fibroblast-specificity of expression (JS_sp_ score in GM00942 primary fibroblasts, excluding GTEx Skin and Heart samples, See Methods).

**Supplementary Figure 9:**
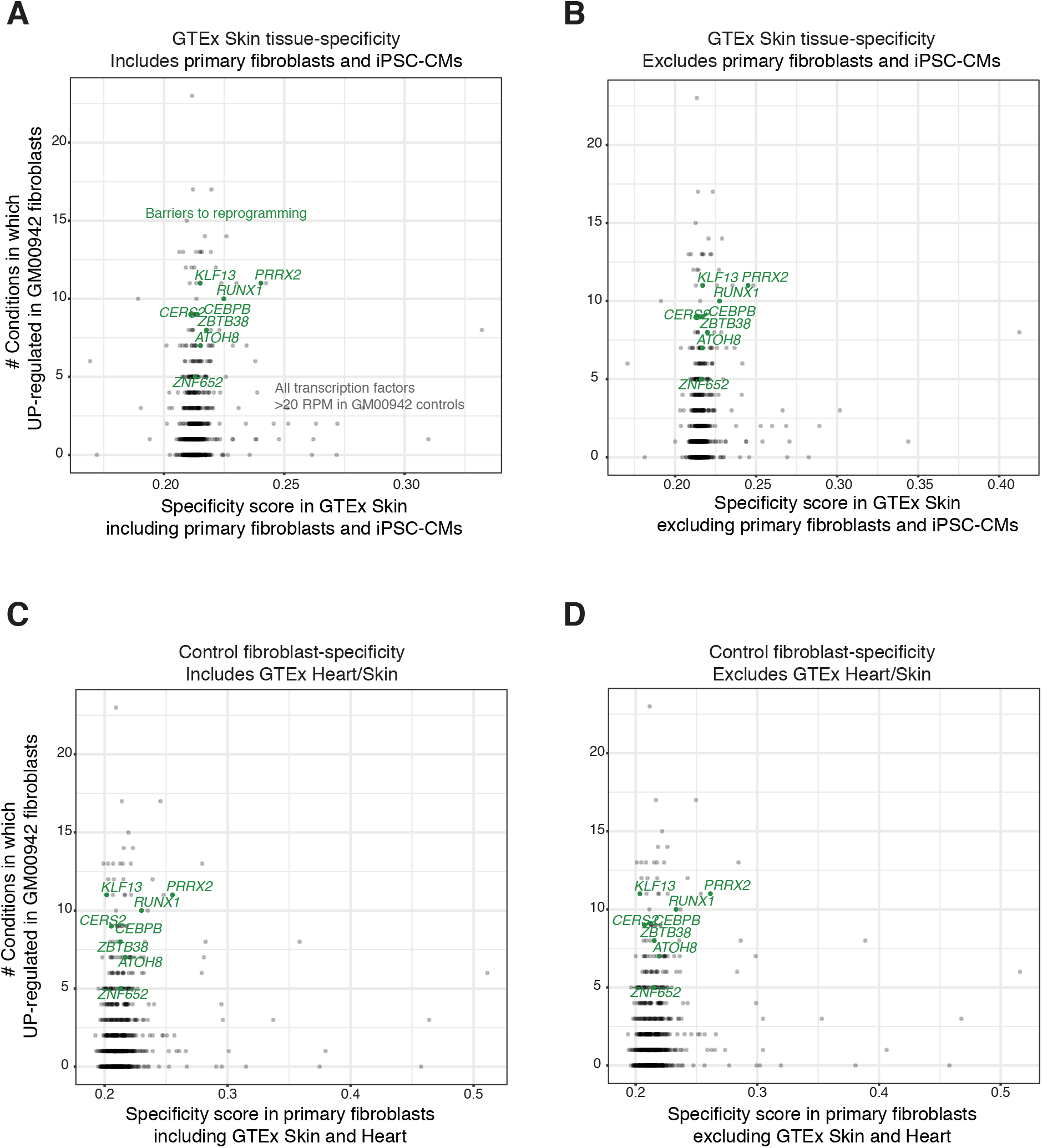
Transcription factor gene expression specificity in fibroblast samples. Expression specificity scores for Skin (A,B) and control GM00942 primary fibroblasts (C,D) are plotted on the x axis (JSsp specificity scores, see Methods). Analyses were repeated on datasets including (A,C) or excluding (B,D) the most similar samples. Responsiveness to perturbation, i.e., the number of conditions in which a gene was expressed higher than in controls, is on the y axis. Panel D also appears in Figure 2.

**Supplementary Figure 10:**
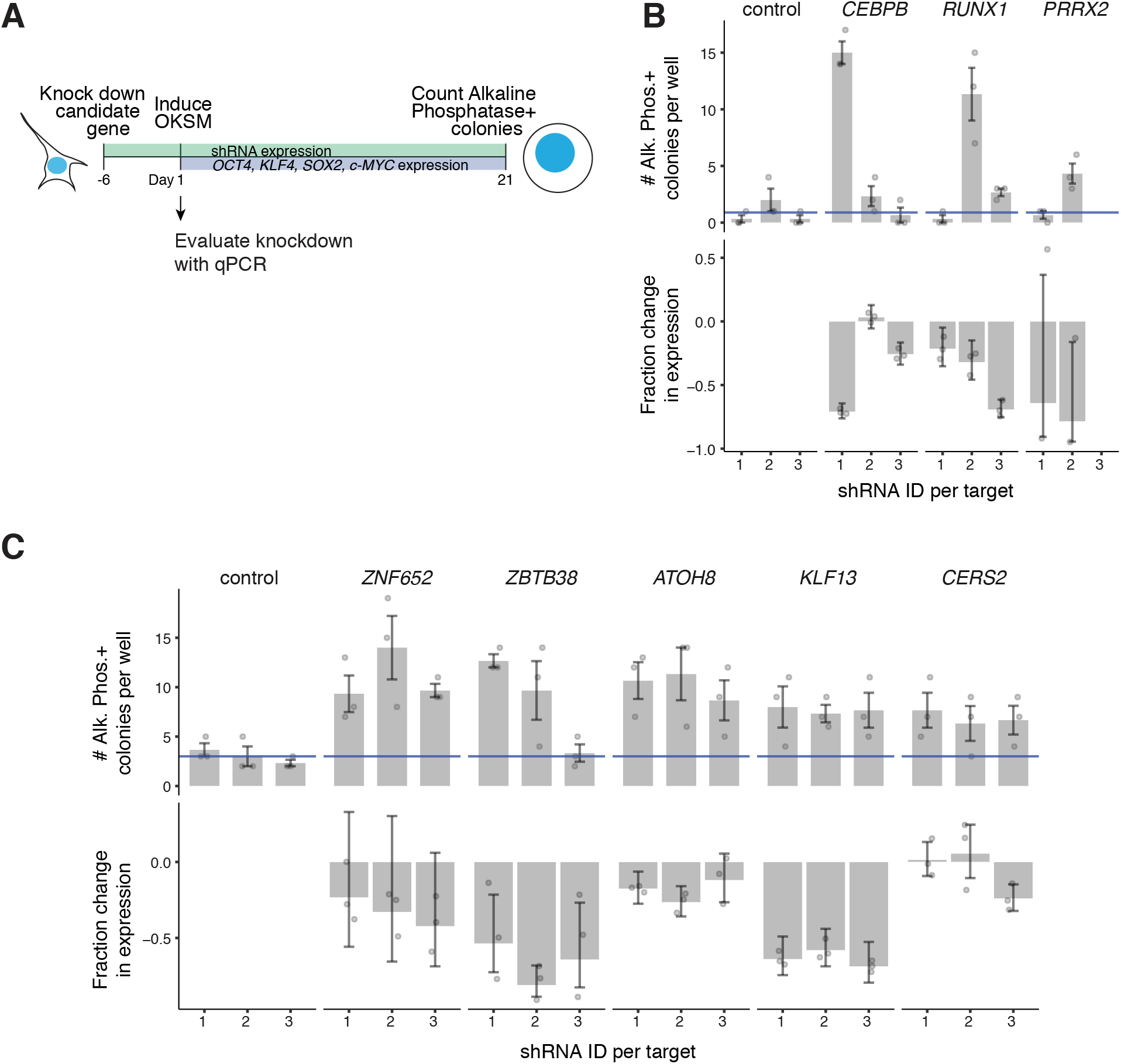
iPSC reprogramming efficiency of hiF-T cells after knockdown of fibroblast-responsive factors. A) Schematic representation of experimental workflow. B, C) Representative replicates. Control shRNAs (1 = empty backbone, 2 = scrambled negative control, 3 = Luciferase-targeting negative control). Controls shown in same row as factors tested are matched per batch. Alkaline Phosphatase-positive colony counts per well counted in triplicate per shRNA. Blue lines correspond to average colony count per well across all control wells in that replicate. Three shRNAs per factor tested. Representative RT-qPCR based assessment of knockdown efficiency.

**Supplementary Figure 11:**
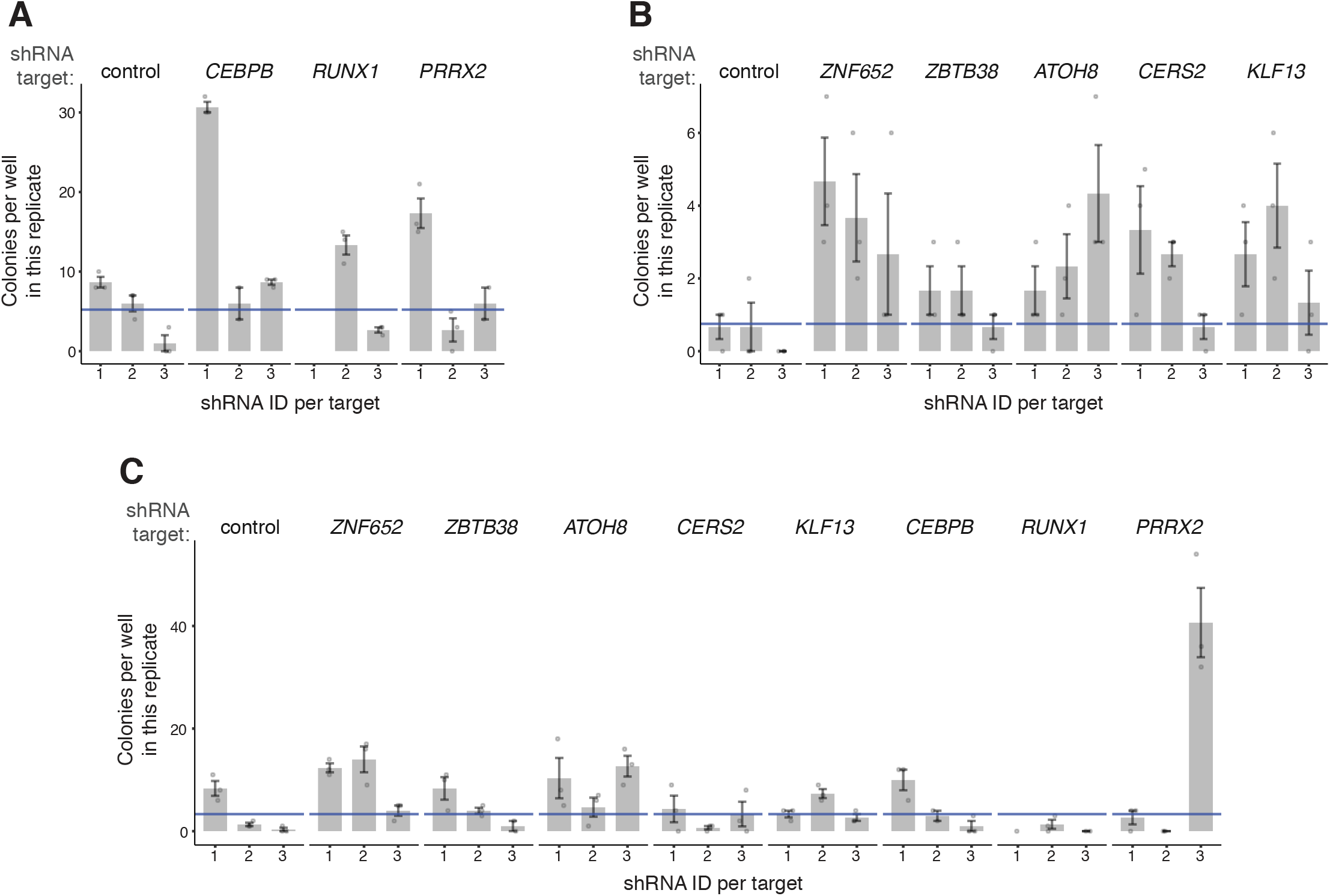
Additional replicates of iPSC reprogramming of hiF-T cells after knockdown of fibroblast-responsive transcription factors. Control shRNAs (1 = empty backbone, 2 = scrambled negative control, 3 = Luciferase-targeting negative control). Controls shown in same graph as factors tested are matched per batch. Alkaline Phosphatase-positive colony counts per well counted in triplicate per shRNA. Three shRNAs per factor tested. Second experimental replicates (A,B), and third experimental replicate (C) for all barriers to fibroblast reprogramming shown in Fig. 2. Blue lines correspond to average colony count per well across all control wells in that replicate. Each experimental replicate was conducted using different vials of cells and different batches of media and feeder mouse embryonic fibroblasts (MEFs) on different days; knockdown samples are compared against controls in the same batch. Reprogramming rate variability was highest in the third replicate for unknown reasons.

**Supplementary Figure 12:**
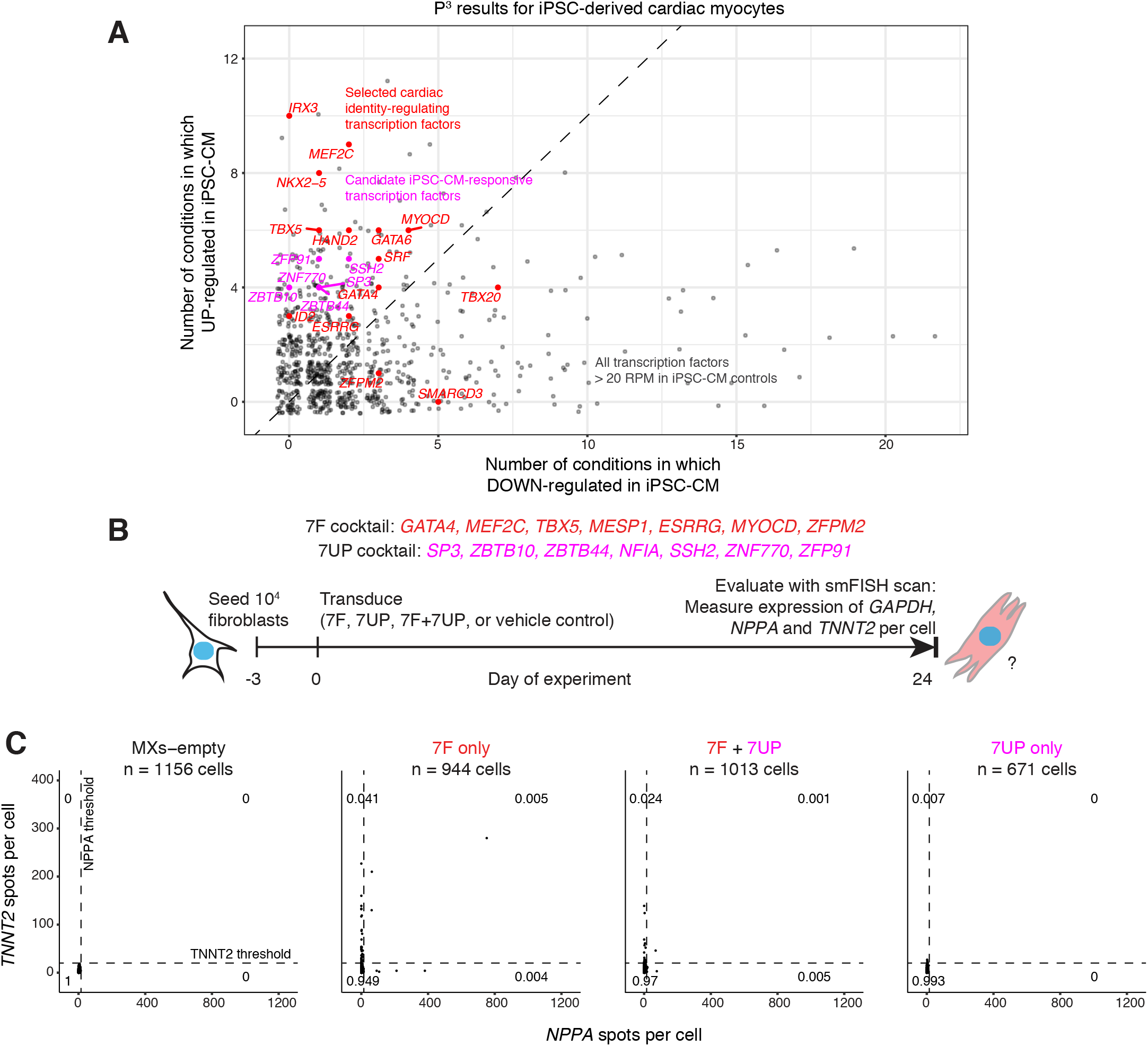
Evaluating cardiac transdifferentiation after overexpression of iPSC-derived cardiac myocyte-responsive transcription factors in fibroblasts. A) Selection of 7 transcription factors found to be frequently up-regulated after perturbation in iPSC-derived cardiac myocytes (iPSC-CM; see Methods). B) Schematic of experimental workflow: iPSC-CM-responsive factors (the 7UP cocktail: *SP3, ZBTB10, ZBTB44, NFIA, SSH2, ZNF770, ZFP91*) were cloned into pMXs retroviral expression vectors, the same system as 7F. 10^4^ immHCF fibroblasts per sample were seeded, transduced with empty pMXs vehicle control (MXs-empty), 7F factors only, 7F + 7UP factors, or 7UP factors only, and cultured in parallel for 24 days (See Methods). C) On day 24, an smFISH scan (1 plane of focus per field of view) was conducted to estimate expression of cardiac myocyte marker genes TNNT2 and NPPA for 671 - 1156 cells per condition. Per cell, we counted the number of TNNT2 and NPPA transcripts per cell, and per-gene quadrant thresholds were chosen arbitrarily to be at least 2 transcripts greater than the maximum expression in MXs-empty condition.

## Discussion

Our model predicted that responsive factors would be important for identity maintenance. We also wanted to test whether responsive factors could in fact help establish cellular identity. Thus, we asked whether the addition of responsive factors to an already established transdifferentiation cocktail would further enhance transdifferentiation. We found that the overexpression of 7UP factors did not enhance 7F-mediated cardiac transdifferentiation, nor was 7UP overexpression alone sufficient for cardiac transdifferentiation. The conclusion would be that the addition of factors to the source cell type that were identified by P^3^ in the target cell type do not help enhance directed changes in cell type. This conclusion fits with our overarching hypothesis, which is that factors identified by P3 are important for identity maintenance (rather than specification). We do note, however, that there are a number of technical limitations that may potentially explain the lack of enhancement. First, each factor is overexpressed by a single virus, so the quantity of virus required to achieve high MOI for all factors could itself be toxic (there is twice as much virus per cell in 7F+7UP as in 7F alone). Additionally, it is possible that the timing of overexpression of 7UP factors relative to the addition of 7F could be important for reprogramming, because as putative identity-maintainers, 7UP factors may increase transdifferentiation efficiency only after a cell has begun the process of identity specification but may interfere with that specification if expressed before 7F has begun to act on a cell. Also, it may be that the stoichiometry of the factors is critical for the enhancement of reprogramming, which is difficult to control experimentally, especially when a large number of factors are added simultaneously.

## References

1. D. Arendt, J. M. Musser, C. V. H. Baker, A. Bergman, C. Cepko, D. H. Erwin, M. Pavlicev, G. Schlosser, S. Widder, M. D. Laubichler, G. P. Wagner, The origin and evolution of cell types. Nat. Rev. Genet. (2016), doi:10.1038/nrg.2016.127.

2. S. A. Morris, P. Cahan, H. Li, A. M. Zhao, A. K. S. Roman, R. A. Shivdasani, J. J. Collins, G. Q. Daley, Dissecting Engineered Cell Types and Enhancing Cell Fate Conversion via CellNet. Cell. 158, 889–902 (2014).

3. P. Cahan, H. Li, S. A. Morris, E. Lummertz da Rocha, G. Q. Daley, J. J. Collins, CellNet: network biology applied to stem cell engineering. Cell. 158, 903–915 (2014).

4. O. J. L. Rackham, J. Firas, H. Fang, M. E. Oates, M. L. Holmes, A. S. Knaupp, FANTOM Consortium, H. Suzuki, C. M. Nefzger, C. O. Daub, J. W. Shin, E. Petretto, A. R. R. Forrest, Y. Hayashizaki, J. M. Polo, J. Gough, A predictive computational framework for direct reprogramming between human cell types. Nat. Genet. (2016), doi:10.1038/ng.3487.

5. Y. Tomaru, R. Hasegawa, T. Suzuki, T. Sato, A. Kubosaki, M. Suzuki, H. Kawaji, A. R. R. Forrest, Y. Hayashizaki, FANTOM Consortium, J. W. Shin, H. Suzuki, A transient disruption of fibroblastic transcriptional regulatory network facilitates trans-differentiation. Nucleic Acids Res. 42, 8905–8913 (2014).

6. C. Chronis, P. Fiziev, B. Papp, S. Butz, G. Bonora, S. Sabri, J. Ernst, K. Plath, Cooperative Binding of Transcription Factors Orchestrates Reprogramming. Cell. 168, 442–459.e20 (2017).

7. H. Weintraub, S. J. Tapscott, R. L. Davis, M. J. Thayer, M. A. Adam, A. B. Lassar, A. D. Miller, Activation of muscle-specific genes in pigment, nerve, fat, liver, and fibroblast cell lines by forced expression of MyoD. Proc. Natl. Acad. Sci. U. S. A. 86, 5434–5438 (1989).

8. E. N. Olson, D. Srivastava, Molecular pathways controlling heart development. Science. 272, 671–676 (1996).

9. J. K. Takeuchi, B. G. Bruneau, Directed transdifferentiation of mouse mesoderm to heart tissue by defined factors. Nature. 459, 708–711 (2009).

10. M. Ieda, J.-D. Fu, P. Delgado-Olguin, V. Vedantham, Y. Hayashi, B. G. Bruneau, D. Srivastava, Direct reprogramming of fibroblasts into functional cardiomyocytes by defined factors. Cell. 142, 375–386 (2010).

11. J.-D. Fu, N. R. Stone, L. Liu, C. I. Spencer, L. Qian, Y. Hayashi, P. Delgado-Olguin, S. Ding, B. G. Bruneau, D. Srivastava, Direct reprogramming of human fibroblasts toward a cardiomyocyte-like state. Stem Cell Reports. 1, 235–247 (2013).

12. L. Qian, E. C. Berry, J.-D. Fu, M. Ieda, D. Srivastava, Reprogramming of mouse fibroblasts into cardiomyocyte-like cells in vitro. Nat. Protoc. 8, 1204–1215 (2013).

13. Y.-J. Nam, K. Song, X. Luo, E. Daniel, K. Lambeth, K. West, J. A. Hill, J. M. DiMaio, L. A. Baker, R. Bassel-Duby, E. N. Olson, Reprogramming of human fibroblasts toward a cardiac fate. Proc. Natl. Acad. Sci. U. S. A. 110, 5588–5593 (2013).

14. T. Vierbuchen, A. Ostermeier, Z. P. Pang, Y. Kokubu, T. C. Südhof, M. Wernig, Direct conversion of fibroblasts to functional neurons by defined factors. Nature. 463, 1035–1041 (2010).

15. J. A. Briggs, V. C. Li, S. Lee, C. J. Woolf, A. Klein, M. W. Kirschner, Mouse embryonic stem cells can differentiate via multiple paths to the same state. bioRxiv (2017), p. 124594.

16. I. A. Mellis, A. Raj, Half dozen of one, six billion of the other: What can small- and large-scale molecular systems biology learn from one another? Genome Res. 25, 1466–1472 (2015).

17. A. B. Keenan, S. L. Jenkins, K. M. Jagodnik, S. Koplev, E. He, D. Torre, Z. Wang, A. B. Dohlman, M. C. Silverstein, A. Lachmann, M. V. Kuleshov, A. Ma’ayan, V. Stathias, R. Terryn, D. Cooper, M. Forlin, A. Koleti, D. Vidovic, C. Chung, S. C. Schürer, J. Vasiliauskas, M. Pilarczyk, B. Shamsaei, M. Fazel, Y. Ren, W. Niu, N. A. Clark, S. White, N. Mahi, L. Zhang, M. Kouril, J. F. Reichard, S. Sivaganesan, M. Medvedovic, J. Meller, R. J. Koch, M. R. Birtwistle, R. Iyengar, E. A. Sobie, E. U. Azeloglu, J. Kaye, J. Osterloh, K. Haston, J. Kalra, S. Finkbiener, J. Li, P. Milani, M. Adam, R. Escalante-Chong, K. Sachs, A. Lenail, D. Ramamoorthy, E. Fraenkel, G. Daigle, U. Hussain, A. Coye, J. Rothstein, D. Sareen, L. Ornelas, M. Banuelos, B. Mandefro, R. Ho, C. N. Svendsen, R. G. Lim, J. Stocksdale, M. S. Casale, T. G. Thompson, J. Wu, L. M. Thompson, V. Dardov, V. Venkatraman, A. Matlock, J. E. Van Eyk, J. D. Jaffe, M. Papanastasiou, A. Subramanian, T. R. Golub, S. D. Erickson, M. Fallahi-Sichani, M. Hafner, N. S. Gray, J.-R. Lin, C. E. Mills, J. L. Muhlich, M. Niepel, C. E. Shamu, E. H. Williams, D. Wrobel, P. K. Sorger, L. M. Heiser, J. W. Gray, J. E. Korkola, G. B. Mills, M. LaBarge, H. S. Feiler, M. A. Dane, E. Bucher, M. Nederlof, D. Sudar, S. Gross, D. F. Kilburn, R. Smith, K. Devlin, R. Margolis, L. Derr, A. Lee, A. Pillai, The Library of Integrated Network-Based Cellular Signatures NIH Program: System-Level Cataloging of Human Cells Response to Perturbations. Cell Syst. 6, 13–24 (2018).

18. B. Szalai, V. Subramanian, C. H. Holland, R. Alföldi, L. G. Puskás, J. Saez-Rodriguez, Signatures of cell death and proliferation in perturbation transcriptomics data-from confounding factor to effective prediction. Nucleic Acids Res. 47, 10010–10026 (2019).

19. M. Niepel, M. Hafner, C. E. Mills, K. Subramanian, E. H. Williams, M. Chung, B. Gaudio, A. M. Barrette, A. D. Stern, B. Hu, J. E. Korkola, LINCS Consortium, J. W. Gray, M. R. Birtwistle, L. M. Heiser, P. K. Sorger, A Multi-center Study on the Reproducibility of Drug-Response Assays in Mammalian Cell Lines. Cell Syst. 9, 35–48.e5 (2019).

20. A. A. Shishkin, G. Giannoukos, A. Kucukural, D. Ciulla, M. Busby, C. Surka, J. Chen, R. P. Bhattacharyya, R. F. Rudy, M. M. Patel, N. Novod, D. T. Hung, A. Gnirke, M. Garber, M. Guttman, J. Livny, Simultaneous generation of many RNA-seq libraries in a single reaction. Nat. Methods. 12, 323–325 (2015).

21. A. Raj, P. van den Bogaard, S. A. Rifkin, A. van Oudenaarden, S. Tyagi, Imaging individual mRNA molecules using multiple singly labeled probes. Nat. Methods. 5, 877–879 (2008).

22. T. M. A. Mohamed, N. R. Stone, E. C. Berry, E. Radzinsky, Y. Huang, K. Pratt, Y.-S. Ang, P. Yu, H. Wang, S. Tang, S. Magnitsky, S. Ding, K. N. Ivey, D. Srivastava, Chemical Enhancement of In Vitro and In Vivo Direct Cardiac Reprogramming. Circulation. 135, 978–995 (2017).

23. K. L. Targoff, S. Colombo, V. George, T. Schell, S.-H. Kim, L. Solnica-Krezel, D. Yelon, Nkx genes are essential for maintenance of ventricular identity. Development. 140, 4203–4213 (2013).

24. Y.-S. Ang, R. N. Rivas, A. J. S. Ribeiro, R. Srivas, J. Rivera, N. R. Stone, K. Pratt, T. M. A. Mohamed, J.-D. Fu, C. I. Spencer, N. D. Tippens, M. Li, A. Narasimha, E. Radzinsky, A. J. Moon-Grady, H. Yu, B. L. Pruitt, M. P. Snyder, D. Srivastava, Disease Model of GATA4 Mutation Reveals Transcription Factor Cooperativity in Human Cardiogenesis. Cell. 167, 1734–1749.e22 (2016).

25. I. S. Kathiriya, K. S. Rao, G. Iacono, W. Patrick Devine, A. P., S. K. Hota, M. H. Lai, B. I. Garay, R. Thomas, Z. Gong, L. K. Wasson, P. Goyal, T. Sukonnik, G. A. Akgun, D. Bernard, B. N. Akerberg, F. Gu, K. Li, W. T. Pu, J. M. Stuart, C. E. Seidman, J. G. Seidman, H. Heyn, B. G. Bruneau, Modeling human TBX5 haploinsufficiency predicts regulatory networks for congenital heart 2 disease. bioRxiv, 835603 (2020).

26. M. Cui, Z. Wang, R. Bassel-Duby, E. N. Olson, Genetic and epigenetic regulation of cardiomyocytes in development, regeneration and disease. Development. 145, dev171983 (2018).

27. GTEx Consortium, Laboratory, Data Analysis &Coordinating Center (LDACC)—Analysis Working Group, Statistical Methods groups—Analysis Working Group, Enhancing GTEx (eGTEx) groups, NIH Common Fund, NIH/NCI, NIH/NHGRI, NIH/NIMH, NIH/NIDA, Biospecimen Collection Source Site—NDRI, Biospecimen Collection Source Site—RPCI, Biospecimen Core Resource—VARI, Brain Bank Repository—University of Miami Brain Endowment Bank, Leidos Biomedical—Project Management, ELSI Study, Genome Browser Data Integration &Visualization—EBI, Genome Browser Data Integration &Visualization—UCSC Genomics Institute, University of California Santa Cruz, Lead analysts:, Laboratory, Data Analysis &Coordinating Center (LDACC):, NIH program management:, Biospecimen collection:, Pathology:, eQTL manuscript working group:, A. Battle, C. D. Brown, B. E. Engelhardt, S. B. Montgomery, Genetic effects on gene expression across human tissues. Nature. 550, 204–213 (2017).

28. M. N. Cabili, C. Trapnell, L. Goff, M. Koziol, B. Tazon-Vega, A. Regev, J. L. Rinn, Integrative annotation of human large intergenic noncoding RNAs reveals global properties and specific subclasses. Genes Dev. 25, 1915–1927 (2011).

29. D. Cacchiarelli, C. Trapnell, M. J. Ziller, M. Soumillon, M. Cesana, R. Karnik, J. Donaghey, Z. D. Smith, S. Ratanasirintrawoot, X. Zhang, S. J. Ho Sui, Z. Wu, V. Akopian, C. A. Gifford, J. Doench, J. L. Rinn, G. Q. Daley, A. Meissner, E. S. Lander, T. S. Mikkelsen, Integrative Analyses of Human Reprogramming Reveal Dynamic Nature of Induced Pluripotency. Cell. 162, 412–424 (2015).

30. K. Takahashi, S. Yamanaka, Induction of pluripotent stem cells from mouse embryonic and adult fibroblast cultures by defined factors. Cell. 126, 663–676 (2006).

31. E. M. Chan, S. Ratanasirintrawoot, I.-H. Park, P. D. Manos, Y.-H. Loh, H. Huo, J. D. Miller, O. Hartung, J. Rho, T. A. Ince, G. Q. Daley, T. M. Schlaeger, Live cell imaging distinguishes bona fide human iPS cells from partially reprogrammed cells. Nat. Biotechnol. 27, 1033–1037 (2009).

32. G. Shi, Y. Jin, Role of Oct4 in maintaining and regaining stem cell pluripotency. Stem Cell Res. Ther. 1, 39 (2010).

33. M. Mall, M. S. Kareta, S. Chanda, H. Ahlenius, N. Perotti, B. Zhou, S. D. Grieder, X. Ge, S. Drake, C. Euong Ang, B. M. Walker, T. Vierbuchen, D. R. Fuentes, P. Brennecke, K. R. Nitta, A. Jolma, L. M. Steinmetz, J. Taipale, T. C. Südhof, M. Wernig, Myt1l safeguards neuronal identity by actively repressing many non-neuronal fates. Nature. 544, 245–249 (2017).

34. R. C. Addis, J. L. Ifkovits, F. Pinto, L. D. Kellam, P. Esteso, S. Rentschler, N. Christoforou, J. A. Epstein, J. D. Gearhart, Optimization of direct fibroblast reprogramming to cardiomyocytes using calcium activity as a functional measure of success. J. Mol. Cell. Cardiol. 60, 97–106 (2013).

35. Y.-J. Nam, C. Lubczyk, M. Bhakta, T. Zang, A. Fernandez-Perez, J. McAnally, R. Bassel-Duby, E. N. Olson, N. V. Munshi, Induction of diverse cardiac cell types by reprogramming fibroblasts with cardiac transcription factors. Development. 141, 4267–4278 (2014).

36. L. Wang, Z. Liu, C. Yin, H. Asfour, O. Chen, Y. Li, N. Bursac, J. Liu, L. Qian, Stoichiometry of Gata4, Mef2c, and Tbx5 influences the efficiency and quality of induced cardiac myocyte reprogramming. Circ. Res. 116, 237–244 (2015).

37. C. Guo, S. A. Morris, Engineering cell identity: establishing new gene regulatory and chromatin landscapes. Curr. Opin. Genet. Dev. 46, 50–57 (2017).

38. H. Zhou, M. G. Morales, H. Hashimoto, M. E. Dickson, K. Song, W. Ye, M. S. Kim, H. Niederstrasser, Z. Wang, B. Chen, B. A. Posner, R. Bassel-Duby, E. N. Olson, ZNF281 enhances cardiac reprogramming by modulating cardiac and inflammatory gene expression. Genes Dev. 31, 1770–1783 (2017).

39. U. Parekh, Y. Wu, D. Zhao, A. Worlikar, N. Shah, K. Zhang, P. Mali, Mapping Cellular Reprogramming via Pooled Overexpression Screens with Paired Fitness and Single-Cell RNA-Sequencing Readout. cels. 0(2018), doi:10.1016/j.cels.2018.10.008.

40. Y. Liu, C. Yu, T. P. Daley, F. Wang, W. S. Cao, S. Bhate, X. Lin, C. Still, H. Liu, D. Zhao, H. Wang, X. S. Xie, S. Ding, W. H. Wong, M. Wernig, L. S. Qi, CRISPR Activation Screens Systematically Identify Factors that Drive Neuronal Fate and Reprogramming. Cell Stem Cell. 0(2018), doi:10.1016/j.stem.2018.09.003.

41. A. Ebrahimi, K. Sevinç, G. Gürhan Sevinç, A. P. Cribbs, M. Philpott, F. Uyulur, T. Morova, J. E. Dunford, S. Göklemez, Ş. Arı, U. Oppermann, T. T. Önder, Bromodomain inhibition of the coactivators CBP/EP300 facilitate cellular reprogramming. Nat. Chem. Biol. 15, 519–528 (2019).

42. M. A. Laflamme, K. Y. Chen, A. V. Naumova, V. Muskheli, J. A. Fugate, S. K. Dupras, H. Reinecke, C. Xu, M. Hassanipour, S. Police, C. O’Sullivan, L. Collins, Y. Chen, E. Minami, E. A. Gill, S. Ueno, C. Yuan, J. Gold, C. E. Murry, Cardiomyocytes derived from human embryonic stem cells in pro-survival factors enhance function of infarcted rat hearts. Nat. Biotechnol. 25, 1015–1024 (2007).

43. W.-Z. Zhu, Y. Xie, K. W. Moyes, J. D. Gold, B. Askari, M. A. Laflamme, Neuregulin/ErbB Signaling Regulates Cardiac Subtype Specification in Differentiating Human Embryonic Stem CellsNovelty and Significance. Circ. Res. 107, 776–786 (2010).

44. W.-Z. Zhu, B. Van Biber, M. A. Laflamme, Methods for the derivation and use of cardiomyocytes from human pluripotent stem cells. Methods Mol. Biol. 767, 419–431 (2011).

45. N. J. Palpant, L. Pabon, M. Roberts, B. Hadland, D. Jones, C. Jones, R. T. Moon, W. L. Ruzzo, I. Bernstein, Y. Zheng, C. E. Murry, Inhibition of β-catenin signaling respecifies anterior-like endothelium into beating human cardiomyocytes. Development. 142, 3198–3209 (2015).

46. S. Shanmughapriya, D. Tomar, Z. Dong, K. J. Slovik, N. Nemani, K. Natarajaseenivasan, E. Carvalho, C. Lu, K. Corrigan, V. N. S. Garikipati, J. Ibetti, S. Rajan, C. Barrero, K. Chuprun, R. Kishore, S. Merali, Y. Tian, W. Yang, M. Madesh, FOXD1-dependent MICU1 expression regulates mitochondrial activity and cell differentiation. Nat. Commun. 9, 3449 (2018).

47. E. W. J. Wallace, J. D. Beggs, Extremely fast and incredibly close: cotranscriptional splicing in budding yeast. RNA. 23, 601–610 (2017).

48. S. M. Shaffer, M. C. Dunagin, S. R. Torborg, E. A. Torre, B. Emert, C. Krepler, M. Beqiri, K. Sproesser, P. A. Brafford, M. Xiao, E. Eggan, I. N. Anastopoulos, C. A. Vargas-Garcia, A. Singh, K. L. Nathanson, M. Herlyn, A. Raj, Rare cell variability and drug-induced reprogramming as a mode of cancer drug resistance. Nature (2017), doi:10.1038/nature22794.

49. G. Yu, L.-G. Wang, Y. Han, Q.-Y. He, clusterProfiler: an R package for comparing biological themes among gene clusters. OMICS. 16, 284–287 (2012).

50. J. M. Vaquerizas, S. K. Kummerfeld, S. A. Teichmann, N. M. Luscombe, A census of human transcription factors: function, expression and evolution. Nat. Rev. Genet. 10, 252–263 (2009).

51. O. Padovan-Merhar, G. P. Nair, A. G. Biaesch, A. Mayer, S. Scarfone, S. W. Foley, A. R. Wu, L. S. Churchman, A. Singh, A. Raj, Single Mammalian Cells Compensate for Differences in Cellular Volume and DNA Copy Number through Independent Global Transcriptional Mechanisms. Mol. Cell, 1–36 (2015).

