## Supplementary figures and images for "Responsiveness to perturbations is a hallmark of transcription factors that maintain cell identity"

### Supplementary Video 1

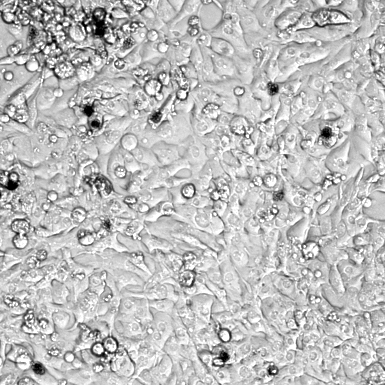

### Supplementary Video 2

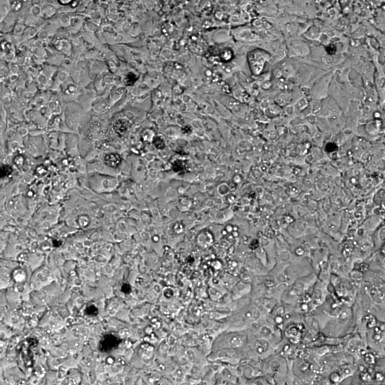
